# Threat of water hyacinth (*Eichhornia crassipes*) on socio-economic and environmental sustainability of Koka and Ziway lakes, Ethiopia

**DOI:** 10.1101/2023.11.10.566637

**Authors:** Esayas Elias Churko, Luxon Nhamo, Munyaradzi Chitakira

## Abstract

Invasive alien plant species cause severe socio-economic and environmental damage. In particular, the water hyacinth (*Eichhornia crassipes*) is an aggressive alien aquatic macrophyte that affects the socio-hydrologic and social environment in many parts of the globe. This study assessed the socio-economic and ecological impacts of the water hyacinth (WH) in Koka and Ziway Lakes in Ethiopia and recommends novel management practices. Purposive sampling design method was used to select households using systematic random sampling. The household sample size was determined with 95% confidence level. Data were collected through key informant interviews, focus group discussion and household surveys, prepared using the Kobo Toolbox which monitors data collectors online. A total of 413 households were sampled and the data were analysed through descriptive statistics and the ANOVA statistical package. At Lake Koka districts that the WH has caused 51% food insecurity by reducing food productivity, and 98.5% health distress through exposure to vector disease. At Lake Ziway districts it caused 81.6% food insecurity and 99.5% health distress. At both lakes, the WH affected the fishing industry by almost 100%. In terms of crop production, maize was significantly affected at Koka, *ᵡ^2^* (1, = 413) = 117.01, p<.001 and cabbage was significantly affected at Ziway, *ᵡ^2^* (1, N= 413) =6.36, p<.001. There was a statistically significant difference in annual income level, age of the household leader, and cost of recovery at household family size, F (9, 623.18) =14.38, p<.001; Wilk’s Λ=.632, partial η2=.14. Therefore, 195 (99.5%) households at Lake Koka and 215 (99.1%) at Lake Ziway illustrated the need for intervention to reduce health impacts and food insecurity. Despite the negative impact, at Lake Koka districts, 86.7% of the plant is used as cattle feed and 28.1% as fertilizer. At Lake Ziway, 42.9% of the plant is used as forage and 39.2%, as a fertilizer.

## Introduction

The Water Hyacinth (WH) (*Eichhornia crassipes*) has severe adverse impacts on aquatic ecosystems at the global scale, making it one of the worst socio-economic and environmental threats (1,2). Previous studies have shown that the distribution and impact of the WH is causing severe economic loss, ecological disruptions, and to humans and the environment health risks (3,4). Many related research findings have shown the negative impacts of WH in the socioeconomic and the environments (5–10).

The global invasive species database indicates that the WH is the worst invasive plant species which is difficult to eradicate as it dominates the water habitat at the expense of native biodiversity (11,12). The WH interferes with the nutrient cycle and food chains as it interacts and alters the structure and function of aquatic ecology, modifying the physical and chemical setting of aquatic systems (13,14). Dense WH diminishes the amount of light reaching hydrophytes submerged in water (15,16). Aquatic fauna, including many fish species, have proper reproduction and growth at the level of dissolved oxygen level more than 4 mgL, but the competition with the WH can cause fish deaths (6,17). Therefore, low dissolved oxygen availability in the water body can negatively affect fish population and density and also challenge other native aquatic species due to high oxygen demand in the water (18). Furthermore, the WH reduces water quantity by increasing evapotranspiration (19,20). From a health perspective, the WH creates the favourable conditions for mosquito breeding and the invasion of snails and other pathogens that cause malaria and bilharzia (9,21,22). A mat of WH over the surface of a water body disturbs tourism, recreation, irrigation, agricultural production, fishing, and transportation on the water body (9,16,23).

Previous studies suggest physical and biological control practice for efficient management of the WH (8). Therefore, since the 19th century, the WH continues to spread from the Amazon basin of South America to the whole world and has since invaded more than 62 countries (24,25). Moreover, the WH has an exceptional reproduction ability and adaptive capacity to invade the new habitat (20,26,27). The plant has a high reproductive capacity and adaptability (8,27–29). The invasive plant reproduces both sexually and asexually, and grows in a faster rate than other perennial water plants, shows high invasion rate, along with embellished expansion globally than many other invasive plant species (8,27,30). The asexual reproduction of the WH is through vegetative reproduction by budding and sexual reproduction is by seeds. Studies show that the WH doubles itself within five to fifteen days in favourable conditions. The optimum growth of WH is at nutrient-rich water body that has a pH value between 6.5 to 8.5 and a water temperature range 28°C to 30°C., and 20 mgL-N, 3 mgL-P, and 53 mgL-K at a salinity < 2% (18,31). Therefore, the ability of expansion of WH is related with its maximum expansion capacity and high adaptability after invading the new environment (8).

In Africa, the first sittings of the WH were recorded in Egypt in 1870’s (28), spreading to many other African countries (4,11). In 1910 it was found in other countries such as such Ethiopia, Kenya, South Africa, Nigeria, Zambia and Zimbabwe (32,33). It was planted intentionally in few places for its beautiful flower and ornamental nature, but in many parts of the world, the weed invaded forcefully causing negative socio-economic impacts on the environment (34). The plant has been destroying native habitats, seriously draining oxygen from aquatic systems and wetlands (26).

In Ethiopia, there are mitigation efforts, research studies and stakeholder activities to control the spread of the WH. The efforts have not been effective due to different reasons and the management or control were also unsuccessful. Some of the reasons were low efficiency of mechanical removal work, shortage of scientifically acceptable dumping site, and limited cooperation by local community members. To enact community participation approach for controlling the WH, studies suggest some methodologies such as empowering opportunities and enticing the locals with potential economic gains (35). Ethiopia has a conservation responsibility for the east African water ecology. Inevitably, the country’s fresh water bodies have seriously been affected by the invasive water plants such as WH, which need continued efforts (9,14,36–39). The nature of the climate of Sub-Saharan Africa and the prevailing weather conditions, particularly hot environments, in the Ethiopian rift valley lakes favour the proliferation and quick expansions of the WH plant in the regions. Additionally, there is eutrophication due to flooding from the farmland that brings artificial fertilizer enter in to the lake water body.

Therefore, the WH control and management needs an integrated approach involving community participation than just mechanical eradication. Assessing WH socioeconomic and environmental impact for the better management of water habitats is vital, particularly, at upper Awash River Basins of the rift valley lakes in Ethiopia. This study was done in the Upper Awash River Basins, in the two-rift valley lakes; Lake Koka, and Lake Ziway of Ethiopia. The aim was to assess the socioeconomic and the environmental impact of WH *(Eichhornia crassipes)* based on the past, present and potential expansion scenarios. The objective was to measure the extent of the social impact of WH in the study districts and characterize the environmental and economic impacts in the invaded locations. The major environmental impairments caused by WH invasion at the riparian community in the study locality were estimated and projected. The social and economic impact of WH in the study area were assessed.

## 2. Materials and methods

**Ethical consideration:** The present study proposal was evaluated by doctoral students’ research proposal evaluation committee of South Africa University (UNISA). The research committee approved the research design and after the research committee approval, the study proposal was presented in the presence of UNISA doctoral students’ research evaluation team, and the academic supervisors. Letter of requesting supports were written from the office of the doctoral students’ research ethics clearance reference number: 2021/CAES_HREC/090, under the Research permission reference number: REC-170616-051. The study locality administration offices after the letter has been offered, approved the research proposal. Then the permission letters were obtained from the respective administrative offices of agriculture, water research chief administrative office and office of environmental protection. The household respondents, FGD participants, and KII persons were fully informed about the purpose of the research. Also, the respondents were informed about how the data obtained from them and would be managed or used as they gave verbal consent. The researcher had checked about informed consent documents obtained from all the participants before the actual interviews were conducted. Finally, key informants and FGDs participant were suggested the places where KII and FGDs should took place. For any further communication, there was a research ethics chairperson of the collage of Environmental study health research ethics Committee.

### 2.1 Description of the study areas

The Awash River Basin, location of the study lakes, is divided in to the Upper, Middle and Lower Valley based on physical and socio-economic factors of the Basin. The Upper, Middle and Lower Valley Basins are parts of the Rift Valleys systems and these sections are seismically active parts of Awash River basin in the Great Rift valley. The current study covers, the two Rift Valley lakes of Ethiopia found in Oromia Regional State at upper Awash River Basin, namely Lake Koka, and Lake Ziway Lake Koka and Lake Ziway are located in the Upper Awash River Basin. The Upper Awash River Basin is located longitude 7°53’7.7"N and 12°14’3.8"N, and latitude 37°58’25"E and 43°18’47.4"E (Fig 1). Lake Koka covers an area of 205km^2^ with the greatest depth of 9m and Lake Ziway covers an area of 442km^2^ and the greatest depth of 8.95m (Fig 1). Lake Koka is located in south central Ethiopia. It is popular with tourists and supports fishing industry, power generation and irrigation. However, it is threatened by sedimentation caused by environmental degradation, and by WH invasion. Lake Ziway is known for its endowment of different bird species and animals such as hippopotamuses. It supports irrigation and fish production. However, in the recent past, deforestation and sedimentation have affected the Lower Awash River Basin, which in turn has also affected the Upper Basin. The Upper Basin has gone through massive degradation due to high population density. Climate and ecological changes in the basin have endangered livelihoods exacerbating the vulnerabilities of the communities living in it.

**Fig 1.**
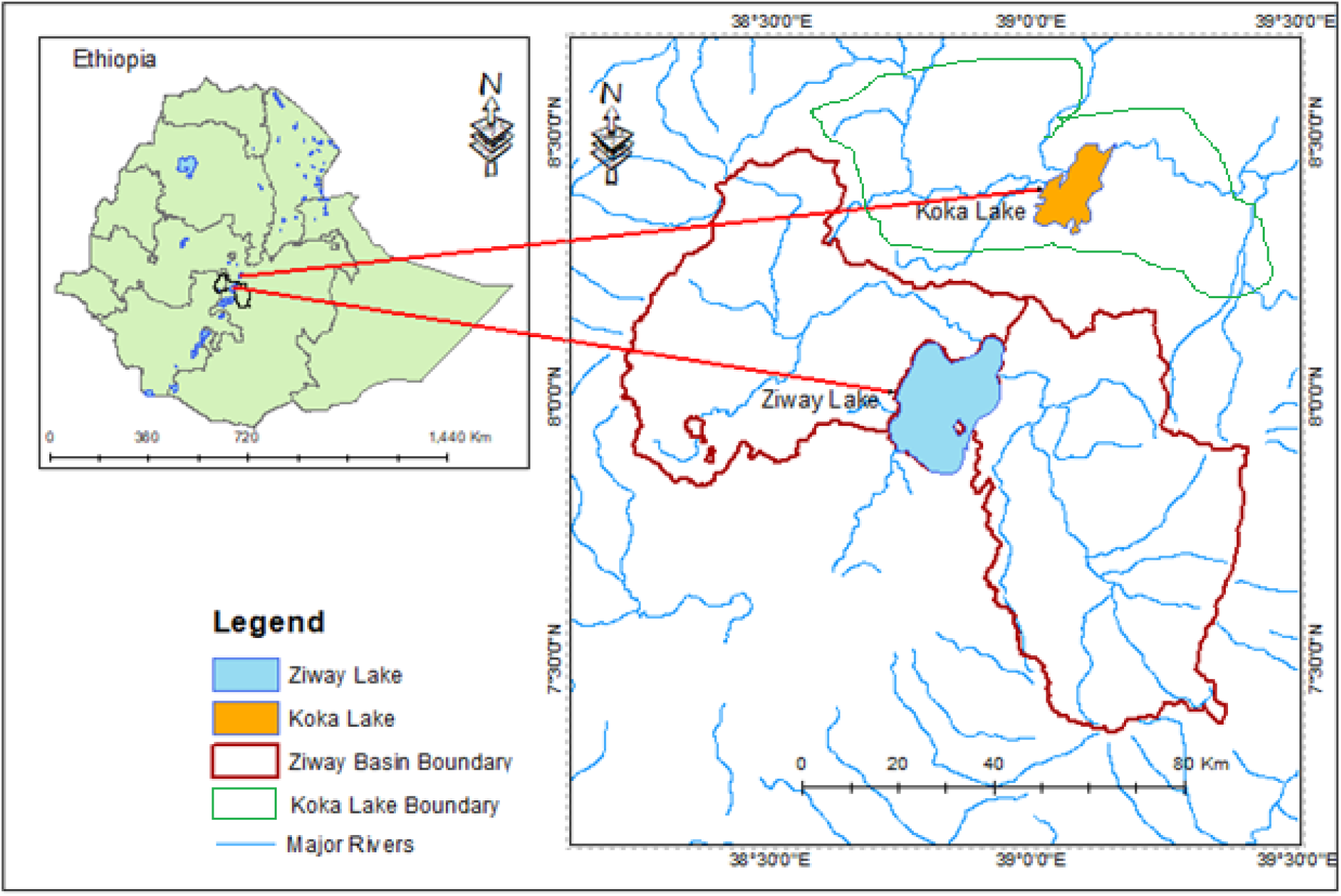
Map of the study areas and sample boarder administrations.

Fig 1. Map of the study areas and sample boarder administrations

### 2.2 Sampling procedure

The study locations were selected by purposive sampling methods to include representative respondents from riparian communities in the two lakes. Two districts from the study lakes, Koka and Ziway, were purposely selected due to high infestation of the WH. Then the sampling of households was done by applying a multi-stage sampling approach as the households had to be close to the lakes. This was done to make sure that only riparian communities that benefit directly from the lakes were selected. The final sampling stage involved sample size determination based on the number of population on the selected districts. After the sampling size determined all the selected household heads completed the questionnaire in the presence of the researcher and research assistants with some technical support while filling the questionnaire.

#### 2.2.1 Sample size determination

The sample size was drawn from riparian communities with a total population of 20,698,000 according to the Ethiopian Census Statistical agency (CSA) (40). In different related researches the sample size is drawn randomly from a total population (41–43). In the present study the total sample size was 20,698,000 and the calculation done as to Taro Yamane formula (44–46). Therefore, from the population to estimate representative sample size, ‘n’ at 95% confidence level was calculated using Equation 1, for the number of persons live in the districts of Lake Koka and Lake Ziway. Accordingly, the calculation of Taro Yamane is presented as follows.

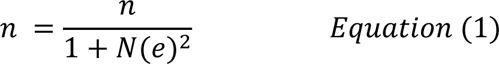

Where: n= the required sample size; N = the population of the area; and e = allowable error (%). Therefore, the sample size will be:

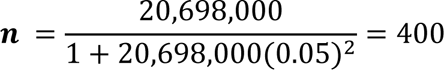

Based on this scenario a total of 400 individuals above 5% contingency or thirteen additional households, totally 413, were participated in the present study. There were a probability that some household might not available for interview or refuse to be respond, thus, thirteen (13) household were added on the sample and the total household become 413. Sample household were methodically selected by picking every n^th^ element of the population, where ‘n’ is the sampling interval. During the selection start up the first household of the sample was selected randomly from the first ‘n’ elements. In this interview the lists of households in each of the respective sample site were obtained from CSA. According to their geographic width and population proportion, fairly all the household representatives were contributed for the interview. A total of four trained assistant’s worked with the researcher from the middle of March up to April 2022, until the whole data collection work had been accomplished.

#### 2.2.2 Sampling design

A total of 413 respondents were sampled, and based on the preliminary household selection the whole sampling was done from each selected households to be included in the selected districts (Table 1). Therefore, data was obtained from the 413 households, key informants (KII) and Focus group discussions (FGD). All the questionnaires focused on the trends of WH spread and its effects on the socioeconomic activity of the community, the communities WH expansion controlling approaches and on the importance of different stakeholder involvement.

**Table 1.** Number and percentage distribution of the total representative sample by study districts.

| Study districts | Number | Percent |
| --- | --- | --- |
| Lake Koka riparian | 196 | 47.5 |
| Lake Ziway riparian | 217 | 52.5 |
|  | 413 | 100% |

In a typical the Ethiopian family the head of the house is the husband (male). However, the woman may head the family in cases where the husband has passed on or a divorced has occurred. Therefore, most of the household heads in this study were males except for a few households which were headed by females. From the total of 413 households of the riparian zone has been participated, the respondents comprised 196(48%) individuals from Lake Koka community and, 217(53%) respondents participated from the Lake Ziway community, as can be seen from the demographics and community status (Table 1).

#### 2.2.3 Data sources

The source of data for the current study were a household questionnaire, KII and FGD. The questions were based on the research objectives and were in line with socioeconomic and environmental protection tools (47). The data were generated in quantitative and qualitative form. For the purpose of triangulation, all the data gathered were backed up by published articles, related periodic and annual reports, and Ethiopian Environmental protection agency short notes on WH, National meteorology, WHO, and other related credible relevant documents of socioeconomic and environmental impacts of WH.

### 2.3 Data collection

As mentioned above data collection were from household survey, FGD and KII. The target population for the household survey were the riparian communities of Lake Koka and Lake Ziway district approximately not more than 5km from each lake. All participants were provided a written consent during the data collection. Data collection was undertaken from April to May 2022. For the household survey, the participants were offered a pre-coded questionnaire of types of items during the survey on socio-economy and environmental impact. The questionnaires were distributed in person to households in the study location. The Kobo Toolbox was used to monitor the data collectors online. The software installed by personal mobile apparatus. All the interviewers were trained operating the software designed for the study on the android version above 6.0. Prior to the actual date interview, an intended household’s survey tested through re-interviewing and crosschecking the response to keep the survey reliability, quality and originality. The key informants participated in this study were the institutional officers such as Environmental officers of the study region, non-governmental organizations work in environmental protection programs, such as environmental and forest conservation specialists, and water development officers. Additionally, community-based organizations head, agriculture department head and the related government agencies and other stakeholders such as wildlife protection officers, chief administrators, and development authority in the location were participated in the interview. Finally, the FGD participants selected by considering the significance of gender equality, religion input, community status, education background and other social values of the community. All the data were backed up and triangulated with the Land use land cover data, condition of vegetation and crops on the fields at the study area, features of the wetlands, nature of pasture lands infested by the WH expansion, and so on.

### 2.4 Data Analysis

By using descriptive statistics such as mean, frequency, and percentage, quantitative data analyzed. Moreover, analysis of variance (ANOVA) was used for different comparisons regarding the WH invasion socioeconomic and environmental impact of riparian community depending on family size at Lake Koka and Lake Ziway. During ANOVA test the data normality was checked using the Shapiro-Wilk Test. showed for both variables were above 0.05, indicating data none deviation from normal distribution. Homogeneity of the data evaluated by Homoscedasticity test showed a value of 0.05 confirming homogeny of the data variance. The analysis was done using the statistical package for social sciences (SPSS) version 27 and Microsoft excel. KII and FGD data taken during the interview were on the audio record and later the Audio - record have been transcribed into English and analyzed according to themes.

## 3. Results

### 3.1 Demographic characteristics and socioeconomic nature of the participants

In the study, the demographics character of participants were assessed, focusing mainly on gender, age-group, marital status, level of education and family size (Table 2). The result reports are discussed as follows. The minimum age range of participants in both gender groups was 18 years, to avoid respondents that were underage. The majority (90.3%) were male while only 40 (9.7%) were female. In terms of marital status, the majority (76%), were married while 24% were either single, widowed or divorced (Table 2). Regarding the education level of the respondents, only 5.3% had no formal education and could neither read nor write while the rest 391 (94.7%) had basic education or had tertiary education. Almost half of the household participants (48.9%) had a total of 3-5 family members, while only 0.9% had a family size above 8 members (Table 2).

**Table 2.** Demographic characteristics of respondents.

| Category and Descriptions |  | Number | Percent |
| --- | --- | --- | --- |
| Gender | Male | 373 | 90.3% |
|  | Female | 40 | 9.7% |
| Age group | 18-24 | 41 | 9.9% |
|  | 25-32 | 118 | 28.6% |
|  | 33-50 | 198 | 47.9% |
|  | >50 | 56 | 13.6% |
| Marital status | Single | 92 | 22.3% |
|  | Married | 314 | 76.0% |
|  | Divorced | 5 | 1.2% |
|  | Widowed | 2 | 0.5% |
| Education level | Cannot read and do not write | 22 | 5.3% |
|  | Can read and write | 71 | 17.2% |
|  | Read and write, basic education | 116 | 28.1% |
|  | High school level | 163 | 39.5% |
|  | Collage level and above, | 41 | 9.9% |
| Family size | ≤2 | 107 | 25.9% |
|  | 3-5 | 202 | 48.9% |
|  | 6-8 | 88 | 21.3% |
|  | ≥9 | 16 | 3.9% |
|  | Total | 413 | 100.0% |

### 3.2 Socioeconomic Impacts of WH

#### 3.2.1 The impact of WH on community’s crop growing

The household participants were offered a pre-coded questionnaire types during the survey on crop production, before and After the WH invasion. As indicated on Table 3, below the survey had a data planned to address each household economic activity and the negative effect of WH on crop production, before and After the WH invasion in the study localities. Crop production is one of the major socio-economic activities in the district. During the data collection the questionnaires were distributed in person to the heads of selected households and the Kobo Toolbox was used to monitor the data collectors online, as mentioned earlier. At actual survey date most of the households were familiar with the technique because, prior to the actual date interview, an intended household’s survey tested through re-interviewing and crosschecking the response to keep the survey reliability, quality and originality. Also, all the interviewers were trained operating the software designed for the study on the android version above 6.0. Majority of the households’ livelihood in the locality were supported based on growing different kinds of crops. The results from the questionnaire survey shown that water shortage for irrigation affected crop production. While as crop production were some of the socioeconomic activities that affected by WH expansion. In the study localities above 90% of the respondents were aware of the invasion of WH in the rift valley lakes of upper Awash River. The findings reveal that WH invasion in the riparian community was becoming a cause for socioeconomic disintegration, crop production failure leading to food shortage and food insecurity (Table 3).

**Table 3.** Crop production trends after WH invasion in the district.

| Crop production After WH invasion |  |  |  |  |  |  |  |
| --- | --- | --- | --- | --- | --- | --- | --- |
| Farm activity and Crops |  | Lake Koka district |  | Lake Ziway district |  | Total Crop production |  |
| Description |  | Before WH | After WH | Before WH | After WH | Before WH | After WH |
| Variables |  | Count/Perc | Count/Perc | Count/Perc | Count/perc | Count/Perc | Count/perc |
| Does WH affect crop production? | Yes | -* | 196(100.0%) | - | 180(82.9%) | - | 376(91.0%) |
|  | No | - | 0(0.0%) | - | 37(17.1%) | - | 37(19.0%) |
| Crops grown<br>in the districts | Carrot |  | 1(0.5%) |  | 4(2.2%) |  | 5(1.3%) |
|  | Tomato | 151(77%) | 75(38.3%) | 180(82.9%) | 89(49.7%) | 331(80.1%) | 164(43.7%) |
|  | Wheat | - | 2(1.0%) | - | 3(1.7%) | - | 5(1.3%) |
|  | Onion | 195(99.5%) | 190(96.9%) | 175(81.1%) | 126(70.4%) | 371(89.8%) | 316(84.3%) |
|  | Maize | 188(95.9%) | 176(89.8%) | 161(74.2%) | 83(46.4%) | 349(84.5%) | 259(69.1%) |
|  | Teff | 144(73.0%) | 40(20.4%) | 137(63.1%) | 4(2.2%) | 281(68.2%) | 44(11.7%) |
|  | G. paper |  | 2(1.0%) |  | 32(17.9%) |  | 34(9.1%) |
|  | Cabbage | 174(88.8%) | 40(20.4%) | 196(90.3%) | 68(38.0%) | 37(89.6%) | 108(28.8%) |
|  | Beans | 54(27.6%) | 0(0.0%) | 81(37.3%) | 28(15.6%) | 135(62%) | 28(7.5%) |
|  | Others |  | 2(1.0%) |  | 5(2.8%) |  | 7(1.9%) |
| Observation:<br>food Stord | Yes | - | 99(50.5%) | - | 216(99.5%) | - | 315(76.3%) |
|  | No | - | 97(49.5%) | - | 1(0.5%) | - | 98(23.7%) |
| Total |  |  | 196(100.0%) |  | 217(100.0%) |  | 413(100.0%) |

\*- = Not applicable or /data was not available

As indicated on Table 3, there were different crops used to grow at Lake Koka and Lake Ziway agricultural habitats. The survey results show that most crops growth and yield after the WH expansion were lower than it was before the WH invasion in the locality. Comparison to the type of crops growing in the districts of the two lakes show that at Lake Koka riparian communities, majority of the households grow vegetables crops more than the beans, cereal crops and other grains (Table 3). As the survey result indicates, types of vegetables grown at Lake Koka districts were Tomato 75 (38.3%), Onion 190 (96.9%), Cabbage 40(20.4%) and among cereals, Maize 176 (89.8%), and Teff 40 (20.4%) (Table 3). Before the WH invasion onion production were (99.5%) at Lake Koka districts, except the onion production, all the other crops production decreased in the districts after the WH invasion. According to the communities’ perception, it was the evidence that WH is affecting food production in their district for two reasons. The first reason was expansion of WH on their crop farm land and the second was the challenge of the irrigation water access blockade due to the dense mat cover the water surface. In different parts of the world there have been reported that natural and manmade factors such as chemicals, floods, and wetland distraction affect normal habitat favouring invasive species dominance by WH (48–50). WH invasion dominate over native water plants, their dens mat create water canal blockade along with the other factors, usually there were a flood line linked to the water bodies that contributes for the expansion of WH, however, negatively affect the growth of normal crops (50–52). Farmers at Lake Ziway districts produce in similar way to the Lake Koka districts. As indicated on Table 3, the production of crops after WH invasion at Lake Ziway was; Tomato 89(49.7%), Onion 126(70.4%), Green Pepper 68(38.0%), and among grains and cereals, Maize 83(46.4%), Teff 32(17.9%), and Beans 28(15.6%), however, the trend shown that the production after WH invasion was lower than before the WH invasion in the district (Table 3). The survey result shown that, except onion production, the total crop production activity in the districts of the two lakes before the WH invasion were higher than after the WH invasion. Therefore, WH invasion has an impact on the lively hood of the community, as it is affecting crop production. For instance, Vegetable crop production is highly dependent on water availability, consequently, in comparison to the two lakes riparian community, households at Lake Koka, who grow vegetable has been affected more than the community at Lake Ziway who grow cereal crops. However, in both circumstances the amount of food production before the WH invasion were higher than after the WH invasion (Table 3).

During the present study, at Lake Koka almost half of the community 99(50.5%) has stored food at visible location and 97(49.5) households didn’t store any food at visible location, which confirms the possibility of insufficient food production in the locality. At the Lake Ziway community, great number of people 216(99.5%) store foods for later consumption, the reason may be at Lake Ziway area the type of crop produced is mainly grains which are easily dried and able to be stored. However, at Lake Koka area vegetable crops are frequently produced, therefore, the impact of WH expansion affected the production of vegetables, which is commonly produced at Lake Koka in the area. Indicating this, among households 196(100.0%) at lake Koka and 180 (82.9%) at Lake Ziway acknowledged their crop production affected after the WH invasion of the locality (Table 3).

The conclusion of current survey results could be that vitality of crop production in the localities has been negatively affected due to the WH invasion. Therefore, the current research findings from the survey result proven that at Lake Koka 195(99.5%) and Lake Ziway 215(99.1%) WH invasion become a serious issue that affecting crop production and causing food insecurity, which needs constant intervention of all stakeholders’ collective action. The participants of FGD at Lake Koka and at Lake Ziway districts explained the challenges of socioeconomic activity after the WH invasion in the area as:

Currently, the WH invasion is a serious problem and a prominent disaster to our livelihood. The groups discussed in detail as follow: Participants of two FGD groups at Lake Koka and three FGD from the Lake Ziway advocated to raise hands for intervention of all stakeholders. Only continued effort can minimize the expansion rate of WH and can increase crop production in our area. The group discussion, particularly at Lake Koka, mentioned human activities such as application of agricultural works enhanced by chemicals, and wetland distraction that cause floods exposes the lake area for increased WH invasion. Additionally, the three FGD team from Lake revealed that in the last five years and the amount of land size owned by the households affected and the level of WH invasion impact on the districts increased. The households owned 0.5 hectare up to 3 hectare farm land near the lakes used to known by the locality as they have built important livelihood in the area, however, most of this people farm lands near the lakes currently invaded by the WH expansion. Lake Koka and Lake Ziway are the two lakes connected by the water banks of Awash River, therefore, thoroughly it the WH transferred to Lake Ziway in the last five years. Two groups of the FGD respondents described that at Lake Ziway WH vastly invaded the lakeshore in the past 5 years. KII’s from environmental protection office of the region and expertise of water research explained quite similar notion on the expansion of WH in the two districts. Currently the WH invasion caused an impact on the community’s crop production and as the result it affected on many households annual income negatively.

Previous studies showed similar results on the possibility of WH expansion that cause negative impact on socioeconomic activities such as loss of Crop and lake water quality impairment (2,53–55). In the present study next to specific crop production trends in the district, the household economic activity and the negative effect of WH invasion on certain crop performance at peak months of WH invasion period in a year, were assessed (Table 4). The data generated from household survey and the objective was to identify peak months of WH infestation that can affect the livelihood of community.

**Table 4.** Perceived peak months of WH infestation that affect livelihood of household.

| Socioeconomic activities |  | Study district |  |  |  |  |  |
| --- | --- | --- | --- | --- | --- | --- | --- |
| Explanation on the study variables |  | Koka Riparian |  | Ziway Riparian |  | Total |  |
|  |  | Count | Percent | Count | Percent | Count | Percent |
| WH spread caused food insecurity | Yes | 196 | 100.0% | 217 | 100.0% | 413 | 100.0% |
|  | No |  |  |  |  |  |  |
| Seasons when WH infestation reaches peak and become critical issue for food production | Jun, July, & August, | 178 | 90.8% | 105 | 48.4% | 283 | 68.5% |
|  | Sept, Oct & November | 16 | 8.2% | 107 | 49.3% | 123 | 29.8% |
|  | Dec, Jan, &, February | 2 | 1.0% | 5 | 2.3% | 7 | 1.7% |

WH invasion has shown different impacts at both dray and wet season’s that are known in the area. As indicated on Table 4, the data were planned to address each household economic activity and the negative effect of WH on crop production, after the peak months of WH infestation that affect livelihood of household in the study localities. In a consecutive three months, Jun, July and August, WH infestation reaches the peak, 283(68.5%), and become critical challenge for the flow of different livelihood activities (Table 4). At the seasons when WH infestation reaches peak and become critical issue for food production, the reason were linked with the amount of rain and flooding in the districts. However, from the month of September up to the month of November 123(29.8%) the WH effect goes down by half and from December up to February it reaches maximum growth and start drying than blooming and multiplication. Socioeconomic and environmental impact of WH invasion that were affecting crop production and causing food insecurity were commonly reported in many socioeconomic studies (53,55,56).

Below in table 5, analysis of loss of each crops grow at the two districts were methodically reported. Chi square analysis was performed on household survey to know types of crops affected across the two lakes due to water hyacinth (Table 5). The objective of survey report on Table 5 was to identify the specific crop that are affected by WH invasion at each districts. Chi square analysis revealed the top five crops (Onion, Maize, *Teff*, Cabbage, Beans) that were affected by WH invasion at the study location by months.

**Table 5.** Chi Square of perceived loss of crop type by study districts.

| Study districts |  |  |  |  |  |  |
| --- | --- | --- | --- | --- | --- | --- |
|  | Koka |  | Ziway |  | Statistics |  |
| Crop |  |  |  |  |  |  |
| Affected | Yes | No | Yes | NO | $\chi^2$ | P |
| Onion | 190 | 6 | 126 | 91 | 86.6 | 0.00 |
| Maize | 176 | 20 | 83 | 134 | 117.01 | 0.00 |
| *Teff* | 40 | 156 | 4 | 213 | 37.28 | 0.00 |
| Cabbage | 40 | 156 | 68 | 149 | 6.36 | 0.01 |
| Beans | 0 | 196 | 28 | 189 | 27.13 | 0.00 |

The result showed that onion production was significantly affected at Lake Koka area, compared to Lake Ziway area, *ᵡ^2^* (1, N= 413) =86.6, p<.001. Similarly, the analysis indicated a maize production was significantly affected at lake Koka, compared to Ziway, *ᵡ^2^* (1, = 413) = 117.01, p<.001. As previous crops loss, production of *Teff* was significantly affected at lake Ziway area, *ᵡ^2^* (1, N= 413) =37.28, p<.001. However, when it comes to loss of cabbages, it was significantly higher at lake Ziway, *ᵡ^2^*(1, N= 413) =6.36, p<.001. Also, the production of beans was significantly affected at lake Ziway because of WH, *ᵡ^2^* (1, N= 413) =27.13, p<.001 (Table 5). However, as indicated by Table 6, the analysis of the loss of crop type by months and lakes shown further differences of WH invasion and crop production in the two lakes. Moreover, the respondents’ perceptions on the level of WH infestation at different months for different crops were assessed and the report indicated on Table 6.

**Table 6.** Chi Square tests of perceived Loss of Crop type by months and Lakes.

| WH invasion |  | WH infestation Months |  |  | Descriptions |  | Statistics |  |
| --- | --- | --- | --- | --- | --- | --- | --- | --- |
| Crops affected | | Jun-Aug | Sep-Nov | Dec-Feb | Total | $\chi^2$ | df | P |
| Onion | Yes | 212 | 101 | 3 | 316 | 6.96 | 2 | 0.03 |
|  | No | 71 | 22 | 4 | 97 |  |  |  |
| *Teff* | Yes | 41 | 3 | 0 | 44 | 13.92 | 2 | 0.00 |
|  | No | 242 | 120 | 7 | 369 |  |  |  |
| Beans | Yes | 23 | 3 | 2 | 28 | 9.74 | 2 | 0.00 |
|  | No | 260 | 120 | 5 | 385 |  |  |  |

Based on the aim of assessing the crop types affected higher than other crops by months and Lakes as a Chi square analysis revealed the top three crops. The analysis shows that the perceived WH impact on production of Onion, *Teff*, and Beans was significantly higher in months of June, July and August than the other months such as September up to Nov nor December up to February (Table 6). The data were obtained from household survey at the study location by months. Chi square analysis proven WH significantly affect the production of onion, *ᵡ^2^* (1, N= 413) =6.96, p<.003, *Teff*, ᵡ*^2^* (1, N= 413) =13.92, p<.001 and beans, *ᵡ^2^* (1, N= 413) =9.74, p<.001 by months of June up to August in a year (Table 6). In the present study in addition to land crop growing trends, water food production were also assessed. The connection between WH invasion and food security is inversely proportional, when constant intervention of all stakeholders on WH control, then there would be proper production of food in the district for the community. WH has a potential to invade wetlands used for farming activities, and can eliminate native biodiversity that has a value for health, food value and productivity of the aquatic system (16,25,57,58).

#### 3.2.2 The impact of WH on fish production

The study revealed an impact of WH invasion, not only by crop growing and crop production but also on fish production, fish quality and accessibility on fishing ground (Table 7). In the study districts, at many households fish was part of family diet and it has a role of playing food security. Therefore, based on the objective of assessing perceived impact of WH invasion on fish productions, a household survey held and the data generated was indicated on Table 7.

**Table 7.**
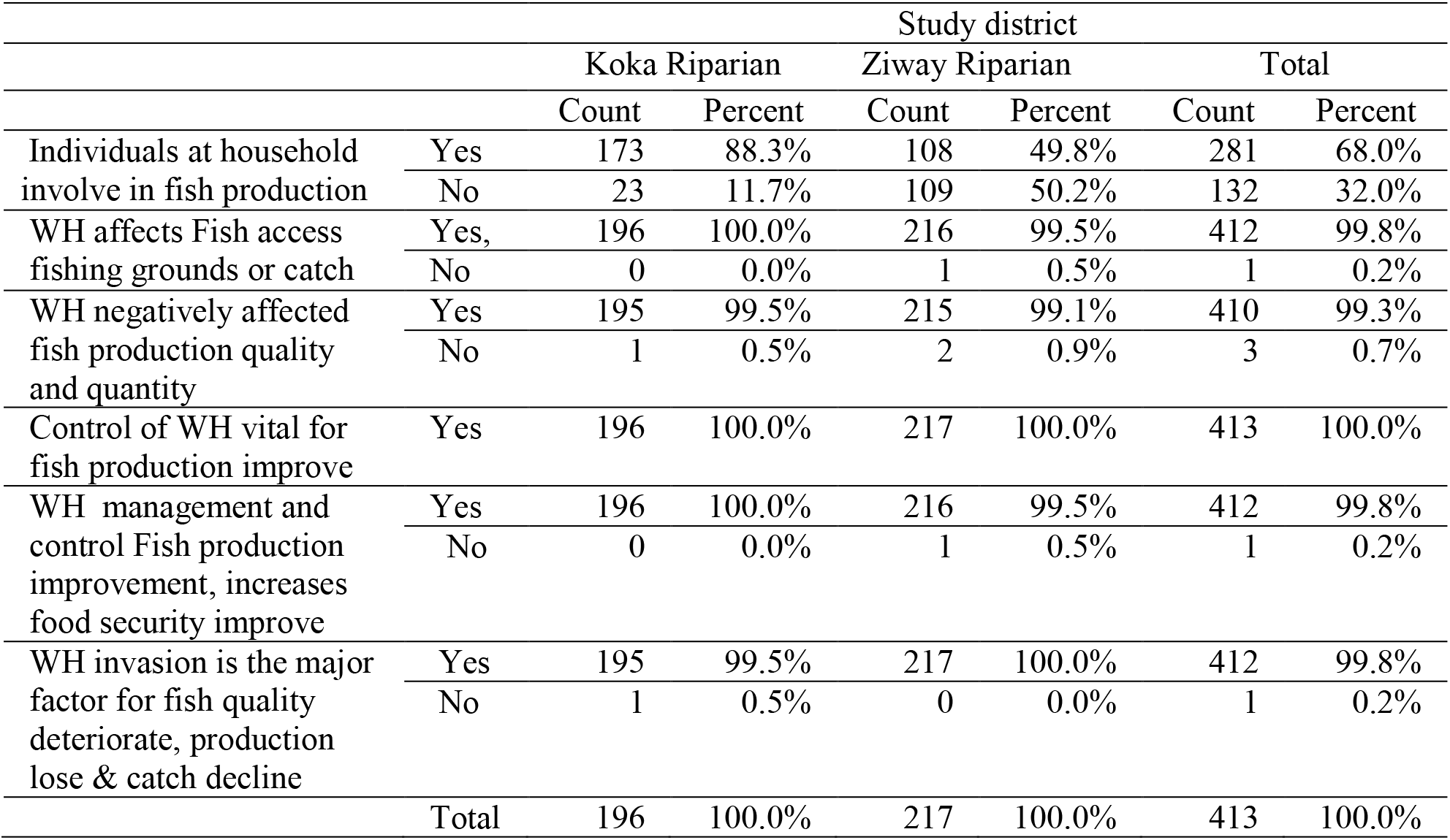
Perceived impact of WH invasion on fish productions.

Quality fish production is part of food security achievement activities, and one of the major water resources utilization (30,59). Fish production can complement food security in a developing countries like Ethiopia. For example; in the study area, 412(99.8%) households confirmed that the better WH management and control, an improved fish production which would increases food security (Table 7). WH invasion of fresh water body causes water quality impairment and apparently causes poor quality on irrigation, fishing difficulties and failure in crop production. The impact of WH invasion causing water quality deterioration in the study districts of Lake Koka and Lake Ziway. Therefore, high infestation of WH means, high impairment fresh water body and low production of fish. In the study location, 412(99.8%) households take a position of WH expansion management failure affected fishing and denied access to the fishing ground (Table 7). Also, the survey results of current study revealed that fish production quality and accessibility is deteriorating and declining in the two lakes due to the water hyacinth spread. At Lake Koka, 195 (99.5%) households and at Lake Ziway 215(99.1%) districts, has shown similar view that WH high infestation means, the fish production is low in quality and few in quantity (Table 7). There was similar findings that supports fish production verses WH occurrence incompatible relationship in the aquatic system, where WH infestation is high, fish production becomes low (17,30,60,61). The research findings on fish production quality lose due to WH attack on water bodies, has shown similar trends at lake Tana Ethiopia, and at elsewhere in south Africa and in China (62–66). According to the majorities of households’ perception, fish production quality deteriorate & catch decline were directly related with the invasion of water hyacinth. The result proven that the expansion of WH affected Fish markets, hinder access to fishing grounds and lowered fish catch at all the study area 412 (99.8%). Generally, the present survey result shown that there were negative impact of WH invasion on fish production as indicated on table 7.

#### 3.2.3 The impact of WH on Annual income of the household

As indicated in the Table 8, an annual income of the total household at Lake Koka and Lake Ziway before and after the WH invasion were assessed. The objective was to compare an annual income of the household before and after WH invasion by number and percent in the study district. The survey data were generated from the household interview (Table 8).

**Table 8.** Annual income of the household before and after WH invasion by number and percent.

| Income (ETB*) | Before WH Invasion |  |  | After WH invasion |  |  |
| --- | --- | --- | --- | --- | --- | --- |
|  | Koka | Ziway | Total | Koka | Ziway | Total |
| 0** | 64(15.5%) | 1(0.2%) | 65(15.7%) | 80(19.4%) | 1(0.2%) | 81(19.6%) |
| <25,000 | 24(5.8%) | 12(2.9%) | 36(8.7%) | 28(6.8%) | 24(5.8%) | 52(12.6%) |
| 25,001-50,000 | 31(7.5%) | 40(9.7%) | 71(17.2%) | 47(11.4%) | 87(21.1%) | 134(32.4%) |
| 50,001-70,000 | 18(4.4%) | 53(12.8%) | 71(17.2%) | 30(7.3%) | 75(18.2%) | 105(25.4%) |
| >70,000 | 59(14.3%) | 111(26.9%) | 170(41.2%) | 11(2.7%) | 30(7.3%) | 41(9.9%) |
| Total | 196(47.5%) | 217(52.5%) | 413(100%) | 196(47.5%) | 217(52.5%) | 413(100%) |

\*\* Not Applicable, \* Ethiopian Birr, 1ETB=0.0178US Dollar

As indicated on table 8, the survey data shows, the five (5) category of yearly income recorded as the household’s respondent. The first category were the household whose Annual income does not show any connection with invasion of WH in the districts. However, the number of these households has increased from 65(15.7%) to 81(19.6%). The data shows those type of households increase in number but it has shown no clear connection with the WH spread in the area. High annual income, above 70,000 ETB per year, was recorded by 170(41.2%) households before the WH invasion at Lake Koka area. It decreased to 41(9.9%) after WH invasion. The lowest annual income, less than 25,000ETB, were recorded by 36(8.7%) households before WH invasion. After invasion it was recorded by more households 52(12.6%) at in the study area. However, the communities’ annual income range is in-between (25,001ETB up to 70,000ETB) in the study location, their annual income increased (Table 8). Generally, in the last decade at many parts of Ethiopia, WH invasion has covered a large area of farm lands and water bodies that weakened crop production and become a reason for food insecurity (57,64,67).

Therefore, the annual income in the study location does not directly related to the spread of WH at Lake Koka or at Lake Ziway districts, except the higher income earning communities (>70,000ETB). The reason is not clearly known at this study but the communities’ income source and socioeconomic activity, however, the total annual income of the locality is not affected by WH invasion and expansion. The results from income and the recovery cost of WH invasion based on age and family size again confirmed the socioeconomic challenges of the community. Household data reported on Table 9 based on the objective of assessing the effect of family size on income, age, and recovery cost of WH invasion.

**Table 9.** The effect of family size on income, age, and recovery cost of WH invasion.

| Multivariate Tests <sup>a</sup> |  |  |  |  |  |  |  |  |  |
| --- | --- | --- | --- | --- | --- | --- | --- | --- | --- |
| Effect |  | Value | F | Hyp. df | Error df | Sig. | Partial Eta Squared | Non cent. Parameter | Observed Power |
| Intercept | Pillai's Trace | .91 | 937.41<sup>b</sup> | 3.00 | 256.00 | .00 | .91 | 2812.21 | 1.00 |
|  | Wilks' Lambda | .08 | 937.41<sup>b</sup> | 3.00 | 256.00 | .00 | .91 | 2812.21 | 1.00 |
|  | Hotelling's Trace | 10.98 | 937.41<sup>b</sup> | 3.00 | 256.00 | .00 | .91 | 2812.21 | 1.00 |
|  | Roy's largest root | 10.98 | 937.41<sup>b</sup> | 3.00 | 256.00 | .00 | .91 | 2812.21 | 1.00 |
| Family size Category | Pillai's Trace | .38 | 12.72 | 9.00 | 774.00 | .00 | .12 | 114.48 | 1.00 |
|  | Wilks' Lambda | **.63** | **14.38** | **9.00** | **623.18** | **.00** | **.14** | **103.11** | **1.00** |
|  | Hotelling's Trace | .55 | 15.68 | 9.00 | 764.00 | .00 | .15 | 141.14 | 1.00 |
|  | Roy's Largest root | .49 | 42.69<sup>c</sup> | 3.00 | 258.00 | .00 | .33 | 128.06 | 1.00 |

There was a statistically significant difference in annual income level, age of the house hold leader, and cost of recovery lost by WH expansion based on household family size, F(9, 623.18)=14.38, p<.001.; Wilk’s Λ=.632, partial η^2^=.14 (Table 9).

As indicated in Table 10, multivariate test result of Wilks’ Lambda supported, the farmland recovery cost after every invasion of WH by family size and age shown, 6-8 households with Std =18556.85 has the income, 128472.43 and this its fair.

**Table 10.** Income and the recovery cost of WH invasion based on age and family size.

|  | Family Size type* | Mean | Std. Deviation | N |
| --- | --- | --- | --- | --- |
| Income | 1.00 <sup>a</sup> | 43972.14 | 21725.23 | 107 |
|  | 2.00 <sup>b</sup> | 39705.56 | 23000.04 | 202 |
|  | 3.00 <sup>c</sup> | 68975.17 | 128472.43 | 88 |
|  | 4.00 <sup>d</sup> | 79585.71 | 60890.88 | 16 |
|  | Total | 49144.33 | 66443.38 | 413 |
| Age | 1.00<sup>a</sup> | 30.25 | 9.07 | 107 |
|  | 2.00<sup>b</sup> | 35.61 | 7.45 | 202 |
|  | 3.00<sup>c</sup> | 45.85 | 10.88 | 88 |
|  | 4.00<sup>d</sup> | 48.43 | 12.18 | 16 |
|  | Total | 37.70 | 10.53 | 413 |
| Recovery cost | 1.00<sup>a</sup> | 22359.13 | 24525.23 | 107 |
|  | 2.00<sup>b</sup> | 8767.32 | 18480.19 | 202 |
|  | 3.00<sup>c</sup> | 11042.74 | 18556.85 | 88 |
|  | 4.00<sup>d</sup> | 4489.71 | 2995.35 | 16 |
|  | Total | 11333.75 | 19778.10 | 413 |

*a. Single household (0) marriage -2 household at a house, b. 3-5 households at a house*
*b. 6-8 households at a house, d. 9-13 households at a house*

### 3.3 Health Impact of WH invasion and Water borne diseases

In the study districts to assess the WH invasion impact on human health by number and percent a household interview were held and the survey result presented on Table 11. WH invasion becoming health determining factors in the study community 409(99.0%). Diseases such as typhoid, malaria, and cholera are some of the diseases that the community 412(99.8%) households suffer endlessly in the districts and the community linked up its prevalence with WH invasion (Table 11).

**Table 11.** WH invasion impact on human health by number and percent.

|  |  | Study Sites |  |  |  |  |  |
| --- | --- | --- | --- | --- | --- | --- | --- |
|  |  | Koka Riparian |  | Ziway Riparian |  | Total |  |
|  |  | Count | Percent | Count | Percent | Count | Percent |
| Currently, water born disease due to WH | Yes | 193 | 98.5% | 216 | 99.5% | 409 | 99.0% |
|  | No | 3 | 1.5% | 1 | 0.5% | 4 | 1.0% |
| Malaria, Typhoid & Cholera occur due WH | Yes | 195 | 99.5% | 217 | 100.0% | 412 | 99.8% |
|  | No | 1 | 0.5% | 0 | 0.0% | 1 | 0.2% |
| Frequently relapsed disease in the districts due to the WH expansion and invasion (Choose one or more) | Malaria | 184 | 93.9% | 189 | 87.1% | 373 | 90.3% |
|  | Typhoid | 9 | 4.6% | 16 | 7.4% | 25 | 6.1% |
|  | Cholera, | 2 | 1.0% | 12 | 5.5% | 14 | 3.4% |
|  | Death of Horse | 1 | 0.5% | 0 | 0.0% | 1 | 0.2% |
| Proclamation needed on WH and regional health | Yes | 186 | 94.9% | 217 | 100.0% | 403 | 97.6% |
|  | No | 10 | 5.1% | 0 | 0.0% | 10 | 2.4% |
| Total |  | 196 | 100.0% | 217 | 100.0% | 413 | 100.0% |

Disease prevention through working on health determining factors needs systematic diseases prevention approaches, and methodology such as changing policy and strategy, new proclamation, and community 403(97.6%) inclusive action plans. WH has been submitted as a human health determinant factor for different reasons. The main reason was its expansion which makes a platform for water borne diseases (21,68,69). The WH invasion effect and its negative aspect on environment and human health contributed for the disease causing vectors a reproductive ground (9,21,22,68). In the current study location the communities usually suffer from malaria, typhoid and cholera. Therefore, 193 (98.5%) at Lake Koka and 216(99.5%) at Lake Ziway, predicted WH infestation were a platform for the reproduction ground of vectors that carry this water born disease (Table 7). In the study districts, malaria was most frequently relapsed at Lake Koka 184(93.9%) and at Lake Ziway, 189(87.1%) than Typhoid and Cholera, however, next to malaria, Typhoid 16(7.4%) and Cholera 12(5.5%) were at Lake Ziway districts than Lake Koka. WH infestation and water born disease has been counted as a major health researches topic for the health of many east African people (4,33,60). Developing new policy and setting proclamation is common at some parts of the world, when there were social-economic insecurity fear and health and safety issue (22,60,70).

The participants of FGD at Lake Koka district explained the challenges of farm land recovery cost at every year rainy seasons when the WH went on invasion and its difficulty on farmers as:

The riparian community of Lake Koka districts have onion and other expensive vegetable crop growing land near the lake shore. The communities have a trend of the manual removal of the WH every year once a while to make free the land from the water hyacinth invasion. During the campaign we get meet up in person and also connected for couple of weeks even a month discussing and spending day and night at work. In the events there were many strange reports from personal experience of different household. We do believe WH has taken away our fertile land, animals eat it, like horses and mules, are die. So far two people get skin disease, and the other two die after working alone for long days and being bit by unknown venom went in to their blood as the bite of water animal in the Lake.

The survey data shows similar results. There was an exceptional case of animal disease that cause a death of Horse at Lake Koka district reported by 1(0.5%) household. This finding was completely new suggestion. There was no similar case on any study such as horse disease verses water hyacinth, so it was the new recommendation by the locality to be a discovery path in the history of animal disease related with WH infestation. From the Lake Koka 186(94.9%) households and from Lake Ziway 217(100.0%) households suggested the importance of specific regional proclamation to prevent health impact of WH (Table 11).

### 3.4 WH invasion and perceptions of the community on Environment protection

Despite of its well-known negative impact, WH has shown unique benefits as reported in many previous researches. Based on this objective WH benefits and perceptions of the society on environment protection assessed and the data garneted presented below on Table 12. Originally, WH plants were used as ornamental plant in many parts of the world, including the Lake Koka dam area of Ethiopia (66,71).

**Table 12.** The benefits of WH and perceptions of the society on environment protection.

|  |  | Study Site |  |  |  |  |  |
| --- | --- | --- | --- | --- | --- | --- | --- |
|  |  | Koka district |  | Ziway district |  | Total |  |
|  |  | Count | Percent | Count | Percent | Count | Percent |
| WH is huge challenge for clean & neat environment | Yes | 194 | 99.0% | 217 | 100.0% | 411 | 99.5% |
|  | No | 2 | 1.0% | 0 | 0.0% | 2 | 0.5% |
| The highest effect of WH expansion over the lake water in your area | The shore | 26 | 13.3% | 215 | 99.1% | 241 | 58.4% |
|  | Near home zone | 26 | 13.3% | 0 | 0.0% | 26 | 6.3% |
|  | animal grazing & other grass lands | 144 | 73.5% | 2 | 0.9% | 146 | 35.4% |
| WH keep fertility of the farm when farmers use it | Yes | 129 | 65.8% | 86 | 39.6% | 215 | 52.1% |
|  | No | 67 | 34.2% | 131 | 60.4% | 198 | 47.9% |
| WH as fertilizer or manure | Yes | 70 | 35.7% | 71 | 32.7% | 141 | 34.1% |
|  | No | 126 | 64.3% | 146 | 67.3% | 272 | 65.9% |
| WH as forage for cattle | Yes | 170 | 86.7% | 93 | 42.9% | 263 | 63.7% |
|  | No | 26 | 13.3% | 124 | 57.1% | 150 | 36.3% |
| Total |  | 196 | 100.0% | 217 | 100.0% | 413 | 100.0% |

Most of the respondents described that WH invaded the two Lakes water body such as the lakeshore in the past 5 years more than it was before. Regarding the response of socio-economic impact of WH, the respondents answer has emphasized on the negative impact of WH in the localities than its benefits. In the current study location, WH expansion affected cattle herding area 144(73.5%), the shore 26(13.3%) and 26(13.3%) place near to residential houses (Table 12). But, at the Lake Ziway around the shore, 215(99.1%) households, reported that WH expansion affected their wetland. In many parts of the earth, WH expansion affected wetlands that serve for different purposes (69,72). Commonly, many East African fresh water habitats, and their wetland areas are fragile in Environmental management practice (60). Watershed management practice and the related environmental rehabilitation works to prevent WH expansion in many parts of the world (73,74). However, in the study locality vast majority of the community, 185(94.4%) households, at the Lake Koka community had been involved in watershed management practice to prevent WH expansion (Table 12). Also, at Lake Ziway, 163(75.1%) households worked in the same way as to the Lake Koka community in an environmental management practice. In the study area, still some communities use WH for the ornamental purpose at Lake Koka about 159(81.1%) households but the rest of the communities, 37(18.9%) has no linking to the plant as ornamental nature of the plant. At Lake Ziway districts, 183(84.3%) households are completely none affiliated to the plants ornamental nature, however, 34(15.7%) households acknowledges the WH ornamental nature. In the present study location, the community has positive attitude for environmental rehabilitation of the water body invaded by the water hyacinth (Table 12). Almost all the community, 410(99.3%) households, agree that despite invading the farm land, if WH have utilized properly it can have unique effect on increasing crop production and thus, at some households the WH applied for soil fertility and the result shown a tendency of improving the soil fertility. However, 273(66.1%) do not suggest the residue of WH usage and application for farm land as fertilizer for the soil, even though, 215(52.1%) households, accept that WH application as fertilizer has been proven by improving their soil fertility (Table 12). Throughout the present data collection phases in all the discussions and respondents’ interview, all the FGD and KII participants respectively, suggested proper controlling of the WH, and strategic management action as a vital recommendation for the community.

## 4. Conclusion

WH expansion has been officially documented as a dangerous invasive plant in many parts of Ethiopia including Lake Koka and Lake Ziway districts. The present study assessed the perceived socioeconomic and environmental effects of WH in the Lake Koka and Lake Ziway. The results shown that the WH spread has affected the important resources of Lake Koka and Lake Ziway that are back bones to the livelihoods of the local community. The effect of WH expansion over the lake water system made irrigation activities very difficult by blocking water access. Its expansion impacted crop production by invading farm fields, and complicating crop production. In the localities annual production capacity affected by the WH expansion and it has been deteriorating due to WH. WH has destructed the lakeshore, where it is the favourable breeding ground for fish, therefore, the local people challenged on accessing Fishing ground. As the result, WH invasion affected seriously the catchability of fish, therefore, fish production decreased and the cost of fishing has increased. This study confirmed invasion of WH is the major push factor for fish production lose along with quality deteriorate & catch decline. Therefore, as to the findings, through WH management and control strategy it’s possible to increase fish production quality and quantity. The WH has also invading and dominating various native grass species that are pleasant for animal feeding, therefore, its spread has increased the occurrence of parasite and disease which are related with the livestock balanced feed supply shortage. Currently, the community frequently suffer from diseases such as malaria, typhoid and cholera in the study locality, because of water born disease regenerations due to WH infestation over the lake water bodies. WH invasion has limited access of water for cattle and home use. In addition to the aforementioned diseases occurrence, recently the new observation of animal disease such as Horse death has come to the community’s attention in the area. Lake Koka and Lake Ziway are among the fresh water bodies of Ethiopia. In terms of environmental management, the wise utilization of fresh water bodies has a great value for the health and safety of natural environment. But, the invasion of WH has infested fresh water of the two Lakes, and as a consequence the plant invasion has harmed the natural vegetation of the country and the local communities. At early years of WH recognition, the communities of Lake Koka and Lake Ziway had a practice of the manual removal campaign for controlling the Expansion. The community devoted for sharing labour cost and financed for the eradication but the participation was not effective and efficient. Therefore, challenges such as decrease of lake water volume, disease encounter, and impairment of water quality, biodiversity decline of some native tree and change of vegetation at the study area are commonly known impacts of WH invasion in the districts. In the present research work, the findings showed WH impact on the nature and environment, its spread effect on environmental protection practices and needs of certain management strategies and importance of stakeholders’ interventions. Therefore, in different Socio-economy and environment factors, Lake Koka and Lake Ziway are affected by WH invasion and expansion. Therefore, this study result revealed to use eradicating methods such as manual, biological or mechanical and more effective, an involvement of all stakeholders is crucial. For further detail studies on WH socioeconomic and environment impact assessment, herein the recommendation is, increasing the scope of study and consenting deferent level of expertise on environmental system analysis and ecosystem service.

## Supporting information

Institutional Ethical Clearance

## Acknowledgments

Our thanks go to all Oromia reginal state administrative leaders for giving permission, for the household survey by helping on identifying the participants of the households, KII, and FGD participants. All the households that have willingly devoted their time to share the challenges in the districts that they have been facing by the invasion of water hyacinth. We would like to acknowledge Oromia regional state agricultural offices, environmental protection, water offices and expertise who were providing information and facilitating administrative linkage with the sampled districts and development agents. The authors acknowledge also UNISA, Jigjiga University and the ministry of Education.

## Author Contributions

Churko, E.E.: Idea conceptualization, proposal development, methodology setup, data curation and analysis, investigation, writing original draft, editing, and revision of the manuscript, visualizing Corresponding. Authors: Nhamo, L. and Chitakira, M.: Supervisors and Co-Supervisor of the project from the start to the end. All authors have read and agreed to the published version of the manuscript.

## Data Availability Statement

All data that supported the success of this study is available from the corresponding author upon rational and reasonable request.

## Supporting information

Questionnaire 1. Household survey questionnaire for socioeconomic and environmental impact assessment of water hyacinth in Lake Koka and Lake Ziway area Questionnaire 2 and 3. KII questionnaire check lists and FGD discussion questionnaire items

