## Supplementary material for "Threat of water hyacinth (*Eichhornia crassipes*) on socio-economic and environmental sustainability of Koka and Ziway lakes, Ethiopia": Institutional Ethical Clearance: 05. Ethics approval ,Human Participant Research, ID 67130100 Esayas Elias Churko.pdf

University of South Africa Research

### FORM 1: 2019

**Research ethics application form for conducting research involving either primary, or a combination of primary and secondary human participant data.**

**For research that involves the explicit use of secondary data, please complete form 2.**

If you have any questions about or require assistance with the completion of this form, please contact your supervisor (master's or doctoral students), or the Research Ethics Administrator of the Research Ethics Committee (011 471 2862 or).

| **IF YOU ANSWER YES TO QUESTION a.1 OR a.2, PLEASE CONTINUE FILLING IN THIS FORM. IF ALL ANWERS ARE NO, CONTACT** | | |
|---|---|---|
| a. The proposed study involves human participants | **YES √** | **NO** |
| a.1 Directly through the collection of primary data | Yes | |
| a.2 Both directly and indirectly through the secondary use of data (If secondary data is the main data source, please complete Form 2, Secondary Data Application) | Yes | |
| b. Collecting personal or confidential information | | No |
| c. UNISA employees, students or data | | No |
| d. Potential conflicts of interest (real or perceived) could arise during the course of the research | | No |

| ***This section is needed for record keeping.*** | | |
|---|---|---|
| **DATE SUBMITTED TO ERC/REC** (*for applicant use) | | |
| **PREVIOUS APPLICATION NUMBER** (*for applicant uses) *(Applicant to indicate a previously allocated application number in case of a resubmission if applicable)* | **Previous Application Number** | **Not applicable** |
| | | **√** |

***This section is for office use only.***

| | |
|---|---|
| **APPLICATION NUMBER** | |
| **DATE PROCESSED (submitted to reviewers)** | |
| **RISK LEVEL *(low, medium or high)*** | |
| **TYPE OF REVIEW (expedited or full committee review)** | |
| **AGENDA DATE** *(For expedited transactions, the agenda date is the date the expedited approval gets reported or ratified at the convened ERC)* | |
| **DECISION OF URERC (approved, referred back, disapproved)** | |
| **DATE OF ISSUING APPROVAL CERTIFICATE OR FEEDBACK LETTER** | |
| **PERIOD FOR WHICH APPROVAL IS VALID (*Valid only as long as approved procedures are followed)** | **From:** **To:** |

**PRIVACY INFORMATION:**

The information you provide on this form is collected for the primary purpose of assessing your research ethics application. This information will also be entered into a database to assist with administration, correspondence, and statistical analyses. These records are accessed by the Unisa Research Ethics Review office bearers and members of relevant committees. Records will be made available to authorised third parties should the need arise such as the National Health Research Ethics Council and Unisa structures (such as University Research Ethics Review Committee). All records will be retained for as long as necessary to achieve the purpose for which it was collected.

**Contents of this application form**

| Item | Page no |
|---|---|
| **RESEARCHER'S DECLARATION** | 3-4 |
| **Section 1 – Researcher(s) details** | **5-7** |
| Section 2 – Risk assessment | 7-11 |
| Section 3 – Details of proposed research | 11 - 12 |
| Section 4 – Proposal summary sheet | 13 - 17 |
| **Section 5 – Data management, analysis and design quality** | **17 - 18** |
| Section 6 – Ethical considerations | 19 - 21 |
| Appendix 1 – Checklist (Separate document) | |
| Appendix 2 – Reviewer form (Separate document) | |

**RESEARCHER'S DECLARATION TO ADHERE TO THE UNISA CODE OF CONDUCT REGARDING THE ETHICS OF THE PROPOSED RESEARCH**

**The declaration should be signed in a separate document and provided to the REC/ERC in a scanned format as part of the application package. PLEASE DO NOT PDF THE APPLICATION FORM BELOW TO ALLOW THE COMMITTEE TO OPEN ATTACHMENTS.**

**By signing below, I ESAYAS ELIAS CHURKO (full name of the main researcher) I declare as follows:**

*Double click on text box selected

| | | |
|---|---|---|
| a) | I completed all the sections of this form that are relevant to the proposed research study according to Appendix A. | ☒ Agree |
| b) | I have not commenced with fieldwork relating to any data collection in relation to the proposed research. | ☒ Agree |
| c) | I have acquainted myself with UNISA's code on research ethics expressed in the UNISA Policy on Research Ethics (link) and the Standard Operating Procedure on Research Ethics Risk Assessment (click on link). I shall fully comply with it. | ☒ Agree |
| d) | I shall conduct the research in an ethically responsible way by demonstrating respect for participants' autonomy, considering a fair risk-benefit analysis and employing fair research procedures. | ☒ Agree |
| e) | I shall conduct the research in strict accordance with the approved proposal. I acknowledge that the approval is valid as long as approved procedures are followed. | ☒ Agree |
| f) | I shall notify the URERC in writing of any adverse events that occur arising from harm experienced by participants. | ☒ Agree |
| g) | I shall notify the URERC in writing if any changes to the research are proposed that may affect any of the study-related risks for the research participants (e.g. methodology, sampling, questionnaire, interview schedule). | ☒ Agree |
| h) | I shall maintain participants' privacy and the confidentiality of records pertaining to the research. | ☒ Agree |
| i) | I shall not use the research and information in a manner that is detrimental to human participants or institutions unless it can be scientifically and ethically justified. | ☒ Agree |
| j) | I shall store research data securely and in accordance with the data management measures indicated in my application/proposal. | ☒ Agree |
| k) | I shall uphold research integrity and refrain from conduct that may taint the integrity of science, including, but not limited to plagiarism, fabrication and falsification of data. | ☒ Agree |
| l) | I shall refrain from the use of human participant data that was collected without a valid research ethics approval for the purpose of this research. | ☒ Agree |
| m) | I shall take the necessary steps to warrant that co-researchers, if applicable, familiarise themselves with the Unisa Policy on Research Ethics. | ☒ N/A<br>☐ Agree |
| n) | I accept the privacy information statement set out on page 2. | ☒ Agree |

**Applicant: Principal Researcher**

Full name in Print:
ESAYAS ELIAS CHURKO

Signature: 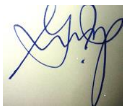

Date signed: April 20/2021

**Approved by supervisor (if applicable):**

To my knowledge the student has addressed all aspects in his/her application for research ethics approval set forth in the University of South Africa's Policy for Research Ethics. I confirm that the form is complete according to Appendix A. I will ensure that the student notifies the committee in writing if any changes to the research are proposed that may affect any of the study-related risks for the research participants such as methodology, sampling, questionnaire, interview schedule, etc. Subsequently, I approve the submission and recommend that approval is granted for the research.

Full name in Print: Luxon Nhamo

Signature: 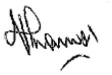

Date signed: 24/05/2021

**Please complete the rest of the form below.**

#### SECTION 1: RESEARCHER'S DETAILS

**This section should be fully completed to aid with the issuing of the clearance certificate and for record keeping.*

| 1.1 | **Details of main researcher (referred to as the applicant)** | | | | |
|---|---|---|---|---|---|
| Title | Full name & Surname | Staff / student no | Department/Unit where you are currently registered or employed | Contact numbers | Email address |
| Mr | Esayas Elias Churko | 67130100 | Environmental science / Environmental management | Mobile: +251911925940, Work: |, |

| | |
|---|---|
| Abridged CV of main researcher explicitly providing evidence of:<br><br>(PLEASE SEE IN ANNEXS) | 1.1.1 Experience relevant to the proposed research<br>1.1.2 Qualifications relevant to the proposed research<br>1.2.3 Publications and other research outputs<br>1.2.4 Research Ethics Training done within the past three years |

| 1.2 | **Supervisor if the application is made by a student** | | | | |
|---|---|---|---|---|---|
| Title | Full Name & Surname | Staff no | Department/Unit where you are employed | Contact numbers | Email address |
| Dr | Luxon Nhamo | 90264126 | Environmental Science | Mobile: 0027 73 166 5859 Work: | |

| | |
|---|---|
| Abridged CV of supervisor explicitly providing evidence of: | 1.2.1 Experience relevant to the proposed research<br>1.2.2 Qualifications relevant to the proposed research<br>1.2.3 Publications and other research outputs relevant to the study<br>1.2.4 Research Ethics Training done within the past three years |

| 1.3 | **Co-supervisor if the application is made by a student***<br>* if applicable | | | | |
|---|---|---|---|---|---|
| Assoc Prof | Chitakira, Munyaradzi | 90227514 | Environmental Science | Mobile: -- Work: 0027 11 471 3220 | |

| | |
|---|---|
| Abridged CV of co-supervisor | 1.3.1 Experience relevant to the proposed research<br><br>1.3.2 Qualifications relevant to the proposed research<br><br>1.3.3 Publications and other research outputs<br><br>1.3.4 Research Ethics Training done within the past three years |

| **1.4** | **Internal and/or External Co-Researcher(s) ***<br>* if applicable | | | |
|---|---|---|---|---|
| Title | Full Name & Surname | Affiliation/ Organisation/Department | Contact numbers | Email |
| | - | - | Mobile:<br>Work: | |
| Abridged CV of co-researcher | | 1.4.1 Experience relevant to the proposed research<br><br>1.4.2 Qualifications relevant to the proposed research<br><br>1.4.3 Publications and other research outputs<br><br>1.4.4 Research Ethics Training done within the past three years | | |

**Please provide information of additional researchers if applicable by inserting additional rows below*

| **1.5** | **Title or provisional title of the research project**<br>*10 - 16 words* | | |
|---|---|---|---|
| **Investigating current and potential expansion, socioeconomic impact and climate smart management approach of Water Hyacinth (Eichhornia Crassipes) in the rift valley lakes of Ethiopia** | | | |
| **1.6** | **Type of application (more than one option may apply)**<br>*Place an 'x' in the box [provide details in the space allocated for comments if applicable]* | | |
| a) Research for non-degree purpose (journal articles; conference presentations, etc.) | | | |
| b) Research for degree purpose | | | (X), |
| c) Identify the qualification for the project (in the case of research for degree purpose) | | | |
| Doctorate in Environmental management | | | |
| d) Collaborative research | | e) Community Engaged Research (CER) | |
| f) Health or Health related research[1]<br>(If you ticked "yes", make sure that you submit this form to a registered Unisa REC/ERC) | | g) Other<br>***Environmental science degree*** | |

[1] Consult the Policy on Research Ethics for a definition of health research.

| | |
|---|---|
| h) Niche Areas *(Unisa researchers only)*<br>i) Knowledge generation and human capital development in response to the needs of South Africa and the African continent<br>ii) The promotion of democracy, human rights and responsible citizenship | X |
| iii) Innovation and capacity building in science and technology | |
| ***iv) Economic and environmental sustainability*** | X |
| v) ODL/ODeL | |
| Comments:<br>Justify why you deem this a CE research project OR collaborative research project OR identify the primary reason for conducting the research if you ticked “Other”. | |

| **1.7** | **Identify the data collection procedures that apply to this research** | **YES (√)** | **NO** |
|---|---|---|---|
| | | *Place an ‘x ‘in the box provided* | |
| a) Survey/questionnaire | | ***‘X ‘*** | |
| b) Focus groups | | ***‘X ‘*** | |
| c) Observations | | ***‘X ‘*** | |
| d) Interviews | | ***‘X ‘*** | |
| e) Documents | | ***‘X ‘*** | |
| f) Other. Please provide details. | | ***Laboratory test and experimental data*** | |

#### SECTION 2 – RISK ASSESSMENT

Complete the Research Ethics Risk Assessment by answering each question below. If you answer **“YES”** to any of the items, the outcome of the risk assessment is considered to vary from a low to high risk level. The UNISA research ethics review system is based on the UNISA Standard Operating Procedure (SOP) for Research Ethics Risk Assessment. If you are an external applicant, a copy of this document can be requested from; internal applicants can click on this link to obtain the document. If you are unsure about the meaning of any of these concepts, please consult your supervisor or project leader.

| **2.1** | **Does your research contribute to knowledge of** | **YES** | **NO** |
|---|---|---|---|
| *Place an ‘x’ in box* ***[if yes, provide details in the space allocated for comments]*** | | | |
| a) | The biological, clinical, psychological or social processes in human beings [social processes refer to those activities, actions, and operations that involve the interaction between people][2] | | X |
| b) | Improved methods for the provision of health services | | X |

[2] *Health* is a state of complete physical, mental and social well-being and not merely the absence of disease or infirmity (Collins English Dictionary)

| | | | |
|---|---|---|---|
| | c) Human pathology | | x |
| | d) Causes of disease | | x |
| | e) Effects of the environment on the human body | | "x " |
| | f) Development or new application of pharmaceuticals, medicines and related substances | | x |
| | g) Development of new applications of health technology referring to machinery or equipment that is used in the provision of health with the exception of medicine[3] | | x |

Comments: If you selected yes to any option above, please describe it in detail here.

**Ethiopian ministry of health; COVID -19 safety equipment & protective protocol for students & researchers:**

The researcher will follow strictly the Ethiopian ministry of health COVID-19 safety equipment and protective protocol. This safety protocol of COVID-19 safety equipment and protective had implemented by the communities in Ethiopian government during the accomplishment of parliamentary election recently. People went for voting and in that case large crowds were at a common place in the election events, where physical distancing measures looks like challenging. Note that, in the protocol the implementation is states wearing face masks, washing hands or sanitizer, keeping social distances and vary at from one local to another. Therefore, when there is any change in the safety equipment and protective measurements, the Ethiopian Ministry of Health provides updates on the status of COVID-19 in the country through its website and social media channels.

| **2.2** | **Does your research include the direct involvement of any of the following groups of participants *(Refer to Section 4 in the SOP)*** | **YES** | **NO** |
|---|---|---|---|
| *Place an 'x' in box [if yes, provide details in the space allocated for comments]* | | | |
| a) Children or young people under the age of 18<br><br>Include the parental consent letter and explain how assent will be obtained in section 6.6 of the application form. | | | "x " |
| b) Persons with disabilities (physical, mental and/or sensory)[4] that could potentially be at risk of harm when participating in this research. | | | "x " |
| c) Persons that might be considered vulnerable, thus finding it difficult to make independent and/or informed decisions for socio, economic, cultural, political and/or medical reasons *(such as the elderly, the dying, unconscious patients, prisoners, those in dependent relationships, women considered to be vulnerable due to pregnancy, victimisation, etc.)* | | | "x " |
| d) Communities that might be considered vulnerable, thus finding it difficult to make independent and informed decisions for socio, economic, cultural, political and/or medical reasons | | | "x " |
| e) UNISA employees, students or alumni<br><br>Indicate that you will apply for permission at the UNISA Research Permission Subcommittee (RPSC) in section 3.1 of the application form to involve any of these participant groups in the proposed research. | | | "x " |

[3] Definition of health research, NHA 61 of 2003, p.8
[4] Describe whether and how proxy or gatekeeper consent will be obtained in section 3.1, 6.1 – 6.3

| | | |
|---|---|---|
| f) Persons who cannot read, speak or understand the language used for the research i.e., English<br><br>Attach the translated data collection instrument(s), interview guide(s), participant information sheet and consent form in the participants' first language, as well as a letter from the language practitioner certifying the credibility of the translated material in Section 4.13. The services of an interpreter may need to be secured for fieldwork activities. | | "x " |
| g) There is a likelihood that a person or definable group will be identified during the research process and it is likely to be of concern. | | X |
| h) Animals | | X |
| i) Other[5]. Please describe. | | "x " |
| Comments: If you selected yes to any option above, please describe it in detail here.<br><br>Through Data collection such as the interview and focus group discussion the respondents will be participated. | | |

| 2.3 | **Does your research involve any of the following types of activity that could potentially place the participants at risk of harm?** | **YES** | **NO** |
|---|---|---|---|
| *Place an 'x' in the box provided [if yes, provide details in the space allocated for comments]* | | | |
| a) Collection, use or processing of personal, identifiable information <u>without</u> the consent of the individual or institution that is in possession of the required information (with the exception of aggregated data or data from official databases in the public domain) | | | "x " |
| b) Collection, use or processing of personal, identifiable information directly from participants <u>with</u> consent (consult the summary of the POPIA on this link). | | | "x " |
| c) Personal, identifiable information to be collected about individuals from available records (e.g. employee records, student records, medical records, etc.) and/or archives | | | "x " |
| d) Personal, identifiable information to be collected outside or transferred outside of South Africa *(if collected from outside you must have consent; if transferred across the border the participant must consent & the country must have adequate privacy laws to protect the personal information)* | | "x " | |
| e) Personal, identifiable information to be shared with third parties for research purposes<br><br>*Attach the confidentiality agreements in Section 6.23 & ensure that prior consent has been obtained from the research participants* | | "x " | |
| f) Participants being exposed to questions which may be experienced as stressful or upsetting, or to procedures which may have unpleasant or harmful side effects | | | "x " |
| g) Participants being required to commit an act which might diminish self-respect or cause them to experience shame, embarrassment, or regret | | | "x " |
| h) Any form of deception of participants, concealment or covert observation | | | "x " |
| i) Examining potentially sensitive or contentious issues that could cause harm to the participants | | | "x " |
| j) Research which may be prejudicial to participants | | | "x " |
| k) Research which may intrude on the rights of third parties or people not directly involved | | | "x " |
| l) Audio-visual recordings of participants which may be of a sensitive or compromising nature (with or without consent) | | | "x " |

[5] Form 1 does not apply to plant, molecular or cell research, animal and environmentally related research.

| | | |
|---|---|---|
| m) Disclosure of the findings of the research could place participants at risk of criminal or civil liability or be damaging to their financial standing, employability, professional or personal relationships | | "x " |
| n) Any form of physically invasive diagnostic, therapeutic or medical procedure such as blood collection, an exercise regime, body measurements or physical examination | | "x " |
| o)*Psychological inventories / scales / tests | | "x " |
| n) Research involving any sensory analysis through the ingestion, smell, taste or feel of food or food related products of any kind. | | "x " |
| p) Other. Please describe | | X |
| Comments: If you selected yes to any option above, please describe it in detail here.<br><br>Data collection site will be from Ethiopia (from riparian residents of the two lakes) which is outside South Africa. | | |

**Please add details on copyright issues related to standardized psychometric tests and registration at the HPSCA of test administrator if test administration is in South Africa or of an equivalent board if administration is non-South African.*

| **2.4** | **Does your research involve any activity that could potentially place the researcher(s) and/or field workers at risk of harm? [if yes, provide details in the space allocated for comments]** | **YES** | **NO** |
|---|---|---|---|
| a) There is a possible risk of physical threat, abuse or psychological trauma as a result of actual or threatened violence or the nature of what is disclosed during the interaction | | | "x " |
| b) There is a possible risk of being in a compromising situation, in which there might be accusations of improper behavior | | | "x " |
| c) There is an increased exposure to risks in everyday life and social interactions, such as working with hazardous materials or sensitive information | | | "x " |
| Comments: If you selected yes to any option above, please describe it in detail here. | | | |

| **2.5** | **Does any of the following apply to your research project?** | YES | NO |
|---|---|---|---|
| *Place an 'x' in the box provided* ***[if yes, provide details in the space allocated for comments]*** | | | |
| a) Participants will be offered inducements or incentives to encourage their involvement in the research | | "x " | |
| b) Participants will incur financial obligations as a result of their participation in the research | | | "x " |
| c) The researcher(s) can anticipate financial gains from involvement in the research (i.e., contract research) | | | "x " |
| d) Any other potential conflict of interests, real or perceived, that could be seen as compromising the researcher(s) professional judgement in carrying out or reporting on the research | | | "x " |
| e) Research will make use of Unisa laboratories | | | "x " |
| f) Research will be funded by UNISA or by an external funding body that could compromise the integrity of the research project | | | "x " |

| |
|---|
| Comments: If you selected yes to any option above, please describe it in detail here.<br><br>During the research there will be 2- 4 volunteer ship research assistants and there will be groups of people from the community that will respond questionnaire may be the team and individuals will be fairly paid some wage as per the agreement to the researcher which may be paid in kind (letter of participation) or cash (per dime as an incentives) for their time and efforts. The payment will be covered from students Bursary or kind sharing of the researcher's pocket money but there is no fund. |

| 2.6 | **Guided by the information above, classify your research project based on the anticipated degree of risk. *[The researcher completes this section. The REC/ERC critically evaluates this benefit-risk analysis to protect participants' rights]*** *Place an 'x' in the box provided* | | | | | |
|---|---|---|---|---|---|---|
| **Category 1**<br>Negligible<br>No to indirect human participant involvement.<br>If you choose this option, stop completing this form and contact | | **Category 2**<br>**Low risk "x "**<br>Direct human participant involvement. The only foreseeable risk of harm is the potential for minor discomfort or inconvenience, thus research that would not pose a risk above the everyday norm. | | **Category 3**<br>**Medium risk**<br>Direct human participant involvement. Research that poses a risk above the everyday norm, including physical, psychological and social risks. Steps can be taken to minimise the likelihood of the event occurring. | | **Category 4**<br>**High risk**<br>Direct human participant involvement.<br>A real or foreseeable risk of harm including physical, psychological and social risk which may lead to a serious adverse event if not managed responsibly. |

(a) Briefly justify your choice/classification

In the study human participant involvement is only by responding questionnaire. Sometimes people fear on giving detailed information, however, the information gathered here will remain confidential. The Questions are related with Water Hyacinth, past, current status and future expansion scenario, its socioeconomic impact and climate smart management activities required in this study area.

(b) Indicate the potential benefits of the study for the research participants and/or communities or other entities. The response will help to learn more about the prospects, challenges and related issues in the area and that will help organizations and communities to facilitate the implementation of Water Hyacinth management in the community. It also gives the government in reviewing and refining the policy, strategy and guiding documents.

(c) Describe the risks relating to the research procedures, which participants, communities or third parties may or will suffer.

There is no risk in actual sense, there may be conditional political escalation some times when people work in natural resource management researches but still here there is no known risk to be mentioned at this moment therefore, this research does not pose any risk to the participants.

| |
|---|
| *This refers to, but is not limited to any participant discomfort, pain/physical or psychological problems/side-effects; persecution, stigmatisation or negative labelling that could arise during the course or as an outcome of the research undertaken.* |
| (d) Indicate how the potential risks of harm will be mitigated by explaining the steps that will be taken to minimise the likelihood of the event occurring (e.g., referral for counselling, debriefing, etc.).<br>When through consenting all the respective local management bodies and discussion with the community leaders the community will become aware that this research may not pose any risk or harm to the participants. |
| (e) Describe the steps to be taken in the case of adverse events or if injury or harm attributable to participation in the study is experienced by the participants, communities or third parties.<br>Offering the permission letter as needed as possible and creating community led awareness about the importance of the study. |
| (f) Describe your arrangements regarding indemnity/compensation for research-related adverse events (if applicable).<br>Not applicable |

#### SECTION 3 – DETAILS OF PROPOSED RESEARCH

Generally, permission to conduct research involving institutions should be obtained prior to field work activities. In case of research involving UNISA employees, students and data, the application form and standard operating procedure can be obtained from **** Application for permission to involve UNISA employees, students and data should be obtained subsequent to ethics clearance.

| **3.1** | **Does your project involve institutions that need to grant permission for research activities?** *Place an 'x' in the box provided* |
|---|---|
| (a) NO | |
| (b) YES | X |

| (c) Name of organisation, state authorities and community advisory board (i.e. UNISA) | (d) Name of person or committee granting permission & contact details (i.e. Research Permission Subcommittee,) | (e) Amount funded/sponsored if applicable | (f) Has permission been granted and is the signed/pro-forma letter attached? *Place an 'x' in the box provided* | | |
|---|---|---|---|---|---|
| | | | YES 'X' | NO | Pending |

(g) Insert the permission letter here or append to the application.
The permission letters are from Oromia Regional office of agriculture and natural resources particularly from irrigation bureau department of natural resource and food security of Ethiopia. In addition to institutions my home University has approved my research work. Please see below:

**Bulchiinsa Mootummaa Naannoo Oromiyaatti**
**Biiroo Qonnaa fi Qabeenya Uummama**
በኦሮሚያ ብሔራዊ ክልላዊ መንግስት የእርሻና ተፈጥሮ ሀብት ቢሮ
**The Regional Government of Oromia**
**Bureau of Agriculture and Natural Resource**

**Guyyaa/ቀን/Date** 27/9/2013
**Lakk./ቁጥር/Ref.No.** [illegible]/1879/[illegible]

**Letter of Authorization**

Mr. **Esayas Elias Churko, ID: 67130100** has requested granting permission for the study to be conducted at lake Ziway and Koka in Ethiopia. We understand this study is for academic purpose and its primary activity will be academic works. The researcher, has submitted the letter of support that states he is currently enrolled for PhD at UNISA College of Agriculture and Environmental science.

The Research intitled: **"*Investigating current and potential expansion, socioeconomic impact and climate smart management approach of Water Hyacinth (Eichhornia Crassipes) in the rift valley lakes of Ethiopia*".** He has been selected the right study area in this title and hereby, our office confirms to you that this research topic is the current top urgent issue and very important subject which needs even high emphases. Therefore, we warmly welcome the researcher and kindly forward for any relevant stakeholders to cooperate for successful accomplishment.

We understand that Mr. **Esayas Elias Churko** will obtain consent for all participants in the study regarding the study site. Mr. Esayas has agreed to work in close contact to our office and he is aware that any data collected for this research work will be kept confidential and will be stored in a secure location per the approved protocol.

Sincerely,

Alému Reggasa Ayane
Deputy Bureau Head &
Irrigation Infrastrucure
Sector Head

Cc:

- Oromia Agriculture And Natural Resource Buureau
- Department of Environmental science at
  UNISA College of Agriculture and Environmental science,

 ፊንፊኔ/አዲስ አበባ

*Deebii Yeroo Kennitan lakkoofsa xalayaa keenyaa caqasaa!*
እባክዎ መልስ ሲሰጡ የደብዳቤያችንን ቁጥር ይጥቀሱ!
**Please quote our Ref. No. While replying!**

**Bulchiinsa Mootummaa Naannoo Oromiyaatti**
**Biiroo Qonnaa fi Qabeenya Uummama**
በኦሮሚያ ብሔራዊ ክልላዊ መንግስት የእርሻና ተፈጥሮ ሀብት ቢሮ
**The Regional Government of Oromia**
**Bureau of Agriculture and Natural Resource**

Guyyaa/ቀን/Date 27/9/2013
Lakk./ቁጥር/Ref.No 5-3/23/02

#### Letter of Authorization

We understand this study is for academic purpose. The researcher, Mr. Esayas Elias Churko,ID:67130100 has requested granting permission for the study to be conducted at lake Ziway and Koka in Ethiopia.

The research intitled**: "Investigating current and Potential expansion, socioeconomic impact and climate smart management approach of water Hyacinth (Eichhorniacrassipes) in the rift valley lakes of Ethiopia."** We welcome Mr. Esayas Elias Churko and kindly confirm you that our office cooperates for successful achievement.

We appreciate permission and consent for all participants in the study as we work in close contact to **Mr. Esayas Elias Churko**and he is awarethat data collected for this research work will be confidential and as well stored in a secure location per the approved protocol.

Sincerely,

Endalkachew Teferi Ligaba
Deputy Head of the Bureau &
Natural Resources and Food
Security Section Head

Cc: - department of Environmental science at

UNISA College of Agriculture and Environmental science,

251 11 371 7455 /011 371 7506 ✉8770 FAX 251 11 371 7488 Finfinnee/Addis Ababa ፊንፊኔ/አዲስ አ[illegible]


*Deebii Yeroo Kennitan lakkoofsa xalayaa keenyaa caqasaa!*
እባክዎ መልስ ሲሰጡ የደብዳቤያችንን ቁጥር ይጥቀሱ!
*Please quote our Ref. No. While replying!*

ጅግጅጋ ዩኒቨርሲቲ
የተፈጥሮና ቀመር ሳይንስ ኮሌጅ
ዲን ፅ/ቤት

**JIGJIGA UNIVERSITY**
College of Natural & Computational
Science Dean Office

Ref .No : ቁጥር --JJU/CNCS/225/13
Date :ቀን :-----17/02/2021

**To Whom it may concern**

**Re: Research Authorization**

This is letter for authority to carry out PhD Thesis research. The research is entitled; "[INVESTIGATING CURRENT AND POTENTIAL EXPANSION, SOCIOECONOMIC IMPACT AND CLIMATE SMART MANAGEMENT APPROACH OF WATER HYACINTH IN THE RIFT VALLEY LAKES OF ETHIOPIA]".

The letter has been issued to our academic staff **Mr. Esayas Elias Churko: ID:67130100** who is attending PhD school at UNISA College of Agriculture and Environmental science. We understand that this study is only for an academic purpose, therefore, it will be conducted maximum within three years.

The student agreed to obtain consent for participants during the study and to provide whenever necessary to any offices the consent linked with the research. Also, upon approval the student will provide the study protocol materials including the approved consent documents in the data collection process whenever necessary.

**Sincerely,**

**Dr. Getu Alemayehu Legesse**

Dr. Getu Alemayehu
College of Natural & Computational Science Vice Dean

College of Natural and Computational Science Dean Office

✉ 1020 ☎0928236192 025-775-5974/5976

(h) If permission is pending, provide an explanation and indicate your planned efforts in obtaining the permission. *Insert a Pro-formal permission letter here or append to the application. Note that the approval for the study may be conditional.*

*Please copy, paste and complete table for additional institutions.*

| | |
|---|---|
| **3.2** | **Are any of the researchers' members of, or do they have any association with the organisations in which you wish to conduct your research?** *Place an 'x' in the box provided* |
| (a) NO | 'x' |
| (b) YES | |

| |
|---|
| (c) If YES, please <u>explain</u> the association clearly in the comment section below. |
| Comments: |

*Place an 'x' in the box provided*

| 3.3 | **Does your research involve collectives and / or communities?**<br>(Group of people sharing social ties, similar interests and a geographic location) |
|---|---|
| (a) NO | |
| (b) YES | **'x'** |
| (c) Please explain what measures you have taken to consult and engage with those communities and / or representative groups regarding your research project below in the comments section. | |
| **Comments:** In this research there will be an interview which requires respondent's participation. Therefore, reaching every one of them will be better after the consensus of the community leaders, and elders in the locality to make formal approach. In Ethiopian culture there is well known practises of discussion habits in group and a small gathering. In this study it is possible to use similar approach. | |

| 3.4 | **Is your project funded or sponsored by any organisation?** |
|---|---|
| (a) NO | 'x' |
| (b) YES | |

| (c) Name of organisation | (d) Name of person or committee granting permission & contact details (i.e., Research Permission Subcommittee,) | (e) Amount funded/sponsored if applicable | (f) Will the identity of any funders be made known to the participants? |
|---|---|---|---|

| 3.5 | **Has this proposal been submitted to another ethics review committee? No**<br>If yes, indicate the name of the institution and the outcome. If previously rejected, provide the reasons. |
|---|---|
| *Insert proof of ethics clearance here | |

| 3.6 | Is this research a sub-study linked to an existing or main study?<br>*Place an 'x' in the box provided* |
|---|---|
| (a) NO | **'x'** |
| (b) YES | |
| Please provide details relevant to the existing or main study in the comments section below | |
| Comments: | |

#### SECTION 4 – PROPOSAL SUMMARY SHEET

**Proposal to be submitted in case of postgraduate student applications, as well as evidence of proposal acceptance by a relevant scientific committee.*

***(*Insert copy of the proposal and the letter of proposal acceptance here)***

| 4.1 | **Introduction, motivation and literature review**<br>*One page (provide a well referenced scientific justification to the study)* |
|---|---|

There are many factors that continue to affect the world's natural resources (Braunisch et al., 2017; Lynch et al., 2010; Thapa et al., 2018). Among these factors; an evasive species expansion dominating over the native species, which has become most harmful factor causing depletion of natural ecosystems is becoming critical issue (Hellmann et al., 2008; Lynch et al., 2010; Thapa et al., 2018; Van Oijstaeijen et al., 2020). Inevitably, every depletion of natural resources effect life on earth. Whatever the extent in the depletion of natural resources, directly or indirectly, it can compound the challenges of resource insecurity and lower the life supporting quality of the earth (Malik, 2007; Wilcock et al., 2008). In Ethiopia, as one of evasive species the challenges of water hyacinth infestation are causing a significant negative impact for the local communities. Due to the plants expansion these riparian community have been facing great economy loss, and social consequences. The plants invasive nature is able to affect millions of riparian zone residents (Firehun et al., 2015; Rinella & Sheley, 2005). These impacts include hazards to human health, shrinkage of lakes and rivers size, wetlands conversion in to dry ground, and decline of native aquatic species biodiversity in availability and composition (Haile, 2019; Hailu et al., 2020; Tewabe et al., 2017). Previous studies from Ethiopia have shown that the water hyacinth affects waterbodies by reducing water quality and quantity (Getnet et al., 2020; Hailu et al., 2020). The plant continued causing many negative impacts in natural environment and health in Ethiopia. Its high expansion capacity and its invasive nature continued posing a serious socioeconomic and environmental problems everywhere in the country where the plant once has been started an expansion (Dersseh et al., 2019; Merga et al., 2020). The plant blocks water ways, causing the blockage of drainage system, weakening power dams, and causing scarcity on pastoral livelihood. Therefore, solving the problems related with this invasive plant and tackling its expansion requires the attention of all stakeholders in the country and beyond. However, there are some documented benefits of the water hyacinth that include, the application in waste water treatment; becoming an excellent source of biomass due to high productivity, and its use as fertilizers (Subash, 2016; Van Oijstaeijen et al., 2020). Despite, few of the benefits, the plant has been affecting a wide range of socioeconomic and environmental impact in the country. This research mainly focuses on the past, the current and future expansion of water hyacinth along with its socioeconomic and environmental impact in the two Rift Valley Lakes, lake Koka or Koka Dam and Lake Ziway of Ethiopia.

| 4.2 | **Research Questions / Hypotheses** |
|---|---|

Research questions.

I. What is the past, the current and potential expansion capacity of water hyacinth in the study area?

II. What is the exact magnitude of the socioeconomic, and environmental problems that water hyacinth causing on the selected Awash River Basin system?

III. Is there any difference on physicochemical variables of the water at the infected and non-infected sites by the Water hyacinth?

IV. What are the benefits, pros and cons for the local community due to Water Hyacinth invasion in the lakes?

V. Which management options is preferable; economically and environmentally, to control further spreading of Water Hyacinth in the upper Rift Valley Basins?

| 4.3 | **Aims and Objectives** |
|---|---|

The aim of this research is to assess the socioeconomic and the environment impact of water hyacinth (Eichhornia crassipes [Mart.] Solms) based on expansion scenarios for integral climate smart management in the Ethiopian Rift Valley lakes.

Specific objectives of the study are to:

- To map the past, the current and future potential expansion of water hyacinth on the upper Awash River Basins; Lake Koka, /Koka Dam, and lake Ziway
- To assess the socioeconomic impact of water hyacinth in the selected study locations
- To identify the physicochemical character of water at a water hyacinth invaded site
- To assess the environmental impact of water hyacinth spreading on the invaded districts
- To provide effective management options for controlling Water Hyacinth expansion

| 4.4 | **Research Paradigm** *Place x in applicable box* |
|---|---|

| | |
|---|---|
| **a) Quantitative** | |
| **b) Qualitative** | |
| **c) Mixed methods** | **"X"** |
| **d) Other**<br>There will be a Lab works and some Observational scientific comments based on systematic speculations will be considered as supplementary by the expertise for further discussion and analysis. | |

Substantiate your choice of paradigm:

Paradigm choice in this research could be validated based on the following reason; the first task will be biodiversity change detection, this is because through GIS and remote sensing the rate of expansion in the past thirty years, right now the current plants' coverage in the area, and the future potential expansion of water hyacinth will be configured on the study location near and around the two selected lakes, which is named as mapping invasive species distribution. The next task will be investigating the socioeconomic impact assessment of water hyacinth followed by the environmental impact of the plant in the study locations. This session of the research paradigm applies Quantitative data collections, along with Qualitative or a Mixed method approach backed up by Observational justification as a supplement by the researcher as expertise.

| 4.5 | **Research Design / Approach / Procedures**<br>*Name & describe the research design you intend to use, e.g., descriptive correlation, case study, grounded theory, etc. If your research will proceed in different phases, describe each phase sequentially.* |
|---|---|

The process of data analysing in this research is by organizing documents in a systematic way to measure the variables under study. The documents will be Qualitative and quantitative data collected in both verbal and written form. After data collection will have been done, all the data will be checked for quality control such as relevancy and reliability then put in to analysis. Both qualitative and quantitative data collected from the respondents will be incorporated as both coded and random data and analysed. Analysis will be based on respondents' answerers such as; questionnaires, expressions, perceptions, behavioural observation, incorporated with photographs, GIS maps, satellite images and records. In addition, data will be analysed by using descriptive statistics representing frequency tables, bar graphs, pie diagrams and percentages. The research design will be descriptive correlation for the

| |
|---|
| three categories of indicator variables such as demographic indicators, accessibility for social infrastructure and economic indicators under the expansion of water hyacinth in the location. The description and correlation will be explained in the research locations, in the communities and observations. In the form of Qualitative and Quantitative data will be Collected for the three indicator and categorized as Discrete and Continuous Variables. In the process of the study the data will be stated as Grouped Data, or any other form of Data Set, and treated for analyses through a SPSS Statistical Software the latest 26 version for analyses. |

| | |
|---|---|
| **4.6** | **Details of the participants of the proposed research project:**<br>**Add additional rows if more than one sampling group is used*<br><br>According the recent Ethiopian official CSA report 2013, the number of persons from the data of Baatu and Adami, districts which is at the vicinity of Ziway and Koka was about 20,698,000, the representative sample size will be 399.99 or about 400. Therefore, when 400 individuals from the study site will be able to participate, then it is possible to consider as the appropriate sample size of participants will have been participated in the study. An individual, respondents' selection will be by using roster random sampling on list-based selection in order to give equal chance for the research participants which are representative sample that reside in the selected study locations. |

| | | | |
|---|---|---|---|
| **4.6.1** | **Describe the participants (in groups) involved in your research project, including the site population, site population size and age category.** | | |
| Identify the participant groups targeted for the research | | Site population size *(How many individuals known to have similar characteristics?)* | Age category of group |
| Group 1* | Household Heads and in a priority list community leader will be given a higher chance to participate than the others who are just a normal house hold heads | $\geq$400 | Above 18 years, since in Ethiopia head of house hold is above 18. |
| Group 2* | Expertise and leaders at managerial at leadership position in different organizations such as agricultural offices, environment and forest conservation offices, water offices, and so on | 36 or more individuals, for at least 6 FGD groups, and minimum 6 officers, from at least 3 organizations | Above 22, that means the average bachelor degree (1st degree) holder may have 22 up to 32 years of age. All the rest may have higher age groups. |
| **4.6.2** | **Explain <u>step by step</u> how you will select participants in each group** *(sampling method, predicted sample size and justification for the sample size).* | | |
| Sampling method | | Sample size | Justify sample size |
| Group 1* | Sampling will be by simple proportional formula: | | Two districts or Woreda / local name for small state in |

| | | | |
|---|---|---|---|
| | $n = \frac{n}{1 + N(e)^2}$ | Population resides near two lakes 20,698,000 | Ethiopian language is called Woreda/ |
| Group 2* | Selection from list by Roster sampling | >400 | woreda |

| | |
|---|---|
| **4.6.3** | **Please specify the inclusion criteria for each participant group.** |
| Group 1* | The population of riparian residents those who live in a woreda around the two lakes |
| Group 2* | Get the Lists of residents and randomly select more or equals to 384 individuals from the roster |

| | |
|---|---|
| **4.6.4** | **Please specify the exclusion criteria for each participant group.** |
| Group 1* | At every house the head of the house hold will be selected as the respondent despite the gender. In Ethiopian context when the house hold among the parents, the male (man) is culturally accepted to give suggestions and comments than the woman at a house hold level or at the village. However, if the man is not alive or is not available by the time then the woman will be selected, finally, if both parents are absent a family member son or daughter above 18 years of age at a house will be selected. The house hold which is in the riparian resident of the two lakes is the target group among the houses and among these houses, the house address located nearer to the lake is selected first than the other houses located side to it. |
| Group 2* | After the name of the residents re listed in the Roster based on the house address near to the lake, the next random selection will be done from the lists. During the selection according to the will of the researcher either odd or even number of names will be selected for the interview. |

| | | |
|---|---|---|
| **4.6.5** | **Describe how much time you require of participants in each group and when the data will be collected/ interviews will take place.** | |
| Time required | | When will data be collected? |
| Group 1* | One time for about 40 minutes | Within a month Tentatively, on June 2021 |
| Group 2* | One time for about 40 minutes | Within two months (Tentatively, July to August) 2021 |

| | |
|---|---|
| **4.6.6** | **Explain how you will obtain the contact details of participants AND provide step-by-step details of how you will recruit them to participate.**<br>If from a public domain source – please identify the source. If from a previously approved database, please confirm how approval was or will be obtained. Attach the approval letter on an official letterhead from the authorised person/committee as an appendix to the application form if you are in possession of it. If not, explain why approval could not be obtained. |
| Group 1* | It will be taken from a previously approved database, please see the confirmation letter of approval which has been obtained from the regional state and others. |
| Group 2* | Lists of people at a woreda near to the two lakes will be taken and re listing will be done for nearby house holds only. |

| | |
|---|---|
| **4.6.7** | **Will any dependent or unequal relationship exist between anyone involved in the recruitment and the participants? [i.e. person in a position of power is recruiting participants which could compromise voluntary participation]** *Place an 'x' in box provided* |
| (a) NO | **x** |
| (b) YES | |

| |
|---|
| Explain if applicable.<br>There is no need for a person in a position of power to involve in recruiting participants rather a person in a position of power can facilitate the research work by understanding the importance of the work. |

| **4.7** | **Collection of data material and procedures** |
|---|---|
| **Indicate which data collection methods will be used.** *Place an 'x' in the box provided* | |

4.7.1 **(a) Questionnaire/survey**

| | |
|---|---|
| YES | "X" |
| NO | |

| | | |
|---|---|---|
| (i) | Self-designed | "X" |
| (ii) | "Borrowed" | |
| (iii) | Adapted | |

| | |
|---|---|
| (iv) | Fully identifiable (name on it) or using a consent form |
| (v) | Potentially identifiable (coded) |
| (vi) | Anonymous (can never be identified) |

| | |
|---|---|
| (vii) | Questionnaire(s)/ survey(s) |
| Insert questionnaire, | |

| | | | |
|---|---|---|---|
| (viii) | If the questionnaire is borrowed, was approval granted by the developers? | Yes | No "X" |
| Insert proof of approval here | | | |
| **(ix)** | **If not, justify why:**<br>**The researcher has been followed Self-designing for the Questionnaire/survey**<br><br>Self-designing for the Questionnaire has been required because the research is original in its content and research design. Also, this research and the surveying are more focuses on searching for the solution as the new finding for the challenges of invasive plants expansion. | | |

**b) Explain how the questionnaire/survey will be administered?**

A pre-test interview will be done with 30 farmers in two sampling areas prior to the actual data collection enabling, the objective is for refinement of the questionnaire. The contents of questionnaire will be a retrospective-focused or an item that assesses the situation of locations before-and after the plant's infestation. That means the items prepared within cross-sectional survey questionnaire to capture the changes due to water hyacinth invasion in the two Lakes. The survey will be administered by 2 up to 4 trained assistance will be used for data collecting after pre-testing of the questionnaire. The questionnaires will be completed by visiting households around their homestead. The survey work of the data collectors will be crosschecked when many interviewers involve to confirm reliability of the collected data at the households being surveyed through re-interviewing.

**c) Specify how the questionnaire/survey will be returned to you to ensure confidentiality.**

On the survey to ensure confidentiality when it administered and return to the researcher after the respondents give the response. This research work is doctoral research and the PhD student is always working in together

| |
|---|
| with trained 2 or 4 assistances and therefore, the student keeps in touch all the time when the questionnaire administered to the respondents and return to the researcher. |

<table>
<tr>
<td>
4.7.2 (a) Interviews<br><br>
YES: x<br>
NO:<br><br>
(i) In-depth: x<br>
(ii) Semi-structured: x<br>
(iii) Unstructured:<br><br>
(iv) Audio taped:<br>
(v) Video taped:
</td>
<td>
(vi) Interview questions/ list of topics attached as addendum to application below<br>
* Insert here<br>
Section 1: Identification<br><br>
Section 2: Demographics and community status<br><br>
Section 3. Socioeconomic impact assessment form<br><br>
3.1 Effects of Water hyacinth expansion on Food security of the community<br><br>
3.2 Effects of water hyacinth expansion on Land use land cover of the community<br><br>
3.3 Effects of water hyacinth invasion on the health of the communities<br><br>
Section 4: The impact of hyacinth expansion on Environment protection<br><br>
4.1 The influence of water hyacinth on protection of Environment & the community<br><br>
4.2 The influence of water hyacinth on agriculture, clean water & farm activities<br><br>
4.3 The influence of water hyacinth expansion from time to time in the districts<br><br>
Section 5: Climate smart management opinion of the community on the plants' invasion,
</td>
</tr>
</table>

<table>
<tr>
<td>
4.7.3 (a) Focus groups<br>
Note: Confidentiality cannot be guaranteed in a group setting – this must be explained as part of the consent process<br>
YES: x<br>
NO:
</td>
<td>
(i) Focus group questions/ list of topics attached as addendum to application<br>
* Insert here<br>
-Demography of respondents,<br>
-Water Hyacinth expansion and its impact such as<br>
I. Los of Social Benefits<br>
II. Los of Health Benefits<br>
III. Los of Economic Benefits<br>
-Water Hyacinth expansion prevention and the plants management related activities done by the villagers<br>
-Community Involvement and expectations,<br><br>
- expectations, actions, and outcomes
</td>
</tr>
</table>

| 4.7.4 **Other** | (ii) Identify, briefly describe each data collection method and insert data collection tools<br>* Insert here<br>Collection of water sample for laboratory experimental test from the plants infested and less or no infested sites near and from the two lakes in the study location. |
|---|---|

| | |
|---|---|
| YES | x |
| NO | |

| **4.8** | Where will the data be collected? If not known, please provide suggested locations. |
|---|---|

Location is already known; GIS**,** remote sensing and satellite images acquisition**;** This will be used to quantify all the past and current expansion image and show the future expansion rate of water hyacinth in the selected area. In all cases, for analysis of the image and results obtained will be validated using field data. At none rainy season or dry period the data collection will be likely in a month of September 2021 up to May 2022 and at any late wet seasons in a year. For quantifying the past distribution of Water Hyacinth, a method is called a new integrative spatial modeling approaches that employ advanced Geographic Information Systems (GIS), advanced remote sensing, along with modeling algorithms (e.g., correlative models).

Therefore, the period of classification is from the past thirty-five years which has been used as one Climate Era Classification. That mean one climate era is 30 years' time or in this study it can be about 35 years range. Since the aim of quantifying the past distribution in the location is to determine the spatiotemporal distribution of water hyacinth in the past thirty-five years range, it can be remarkable point for time to use for classification. The data obtained in this way, will be triangulated by an intensive literature review and key informant interview regarding the past distribution as known by the district residents. Likewise the current and the future potential distribution will be computed.

**Primary data**; All Household survey, Focus Group Discussions (FGDs) and Key Informant Interviews (KIIs) will be used for primary data collection based to the research design from the location where the water hyacinth plant invasion observed at the upper awash river. The data will be collected from the selected household survey, discussion with the Focus Group (FGDs) and by interview from Key Informant Interviews (KIIs) at riparian residents of the two lakes mentioned above. The data generated will be both quantitative and qualitative type. Here a qualitative data to augment a quantitative data (survey) output. The use of qualitative approach relies on phenomenology as an approach to qualitative study. This approach emphasizes on socioeconomic survey of the community's subjective experiences and interpretations. And also, for environmental impact assessment of the plants in the study locations. Qualitative method provides an opportunity to uncover unexpected results or unforeseen contextual factors.

**Data Triangulation**: Data that helps for triangulation will be gathered from the annual reports of EEPA, national meteorology, periodically published magazines of the residents, Awash River Basin Tourism Development Office, WHO, and Awash River Public Water Enterprise, WFP, and many more credible relevant documents. The data from reports of these institutions will include the number of people engaged in the weeding campaign, and the area covered by water hyacinth and the people general perception about the plant expansion. Additional data on Land use land cover data such as the size of crop fields and pasture lands infested by the weed, the effect of the plant on water transportation and hydroelectric power will be collected. Likewise, any secondary relevant data about socioeconomic impacts of water hyacinth will be collected from electronic and hard copy literature sources, and Journal review articles. In the aforementioned approach both primary and secondary data will be collected; these methods are believed as a sufficient technique for the socioeconomic impact assessment and likewise the data collection follows similar steps for Environmental impact assessment. Regarding secondary data, all the related article and intensive literature review will be collected from

| |
|---|
| Ethiopian Environmental protection agency to assess environment impact of water hyacinth in the study area.<br><br>**Water sample**: In-situ measurement of water physicochemical character that help to determine water quality are: the water pH, Temperature, electrical conductivity, Nutrients, TDS, Turbidity, BOD, dissolved oxygen, heavy metals and flow velocity. Water smell: tested through observation and on its smell, weather the wastewater had an objection to the noise or not. |

| | |
|---|---|
| **4.9** | **By whom will the data be collected? (Researcher/field workers/community members)?**<br>Explain any measures that you will take to prepare yourself/field workers/community members to optimise data collection activities. Field workers/community members are required to sign a confidentiality agreement form. |
| | The work will be done by PhD researcher and with only four trained assistants who will work with the researcher for two months possibly from June up to August 2021. |
| * Insert confidentially agreement for fieldworkers/community members here (Please, the Annex parts) | |

| | | |
|---|---|---|
| **4.10** | **Will participants be subjected to any form of intervention (manipulation of the participant or the participants' environment)?** *Place an 'x' in the box provided* | |
| NO | X | YES |
| Please explain the intervention in full. | | |

| | | |
|---|---|---|
| **4.11** | **Does the research involve participants who have specific cultural needs, protocol requirements or/and specific consent arrangements?** *Place an 'x' in the box provided* | |
| NO | X | YES |
| Please explain the intervention in full. | | |

| | | |
|---|---|---|
| **4.12** | **Will you require the use of a translator or will you use documentation translated into a language other than English?** *Place an 'x' in the box provided* | |
| NO | X | YES |
| Describe how the translator will be used.<br>*Attach the translated data collection instrument(s), interview guide(s), participant information sheet and consent form in the participants' first language, as well as a letter from the language practitioner certifying the credibility of the translated material.* | | |

| | |
|---|---|
| **4.13** | **Is there a dependent or unequal relationship between any person collecting the data (e.g. researcher) and the participant?** *Place an 'x' in the box provided* |

| NO | X | YES |
|---|---|---|

Please give details and explain the measures taken to manage this situation.

**NA (Does not apply)**

| **4.14** | **Does your research project involve the collection and analysis of documents or secondary data?** *Place an 'x' in the box provided* |
|---|---|

| YES | NO |
|---|---|
| **X** | |

a) **Please explain the sampling method of the relevant categories of documents and the predicted sample size, followed by a justification for sample size (add more rows if necessary)**

| | Sampling method | Sample size | Justify sample size |
|---|---|---|---|
| Data set/document 1* | Secondary data from the weather forecasting and meteorology offices, | The past 35 years data on the rainfall and temperature | Data in the range of 30 -35 years in together at the selected woreda in the justified location. NB: One climate session. |
| Data set/document 2* | Credible data base from water offices, Ethiopian Environmental Protection office, and from agricultural offices, and any relevant secondary data regarding the water hyacinth expansion and intervention has done so far in the study location. | Any data in the aforementioned research period, can be applied as sample size to use the data in this research. | Data produced in the years that are included will be included as relevant data for this research. There is no special reason for justification at this moment regarding detail justification of the data but years of the data base is enough justification. |

b) **Describe the <u>conditions</u> under which the data was collected initially and the <u>reasons why</u> it was collected. If applicable, describe the <u>number of participants</u> and <u>demographics</u> applicable to the secondary data analysis.**

NA

c) **Was ethical clearance granted for the original data gathering phase by this or by another research ethics committee if appropriate?**

Yes

#### SECTION 5: DATA MANAGEMENT, ANALYSIS AND DESIGN QUALITY

| **5.1** | **<u>Describe</u> the data analysis method that you will use (qualitative data analysis method or quantitative statistical procedures).** |
|---|---|

The two dependent variables in this study will be economic indicator with its items and selected social infrastructure indicators which has a direct contribution to the objectives under this study. While as the other two independent variables are water hyacinth expansion in the location and some selected demographic indicator items that will have a direct impact in the future findings. In the study a set of statistical methods that can apply for the estimation of relationships between these variables. The data generation will be based on the structured questionnaire which responded by the participants. After collection of the data, it will be subjected for analyses using different relevant statical methodologies such as descriptive statistics; mainly in percentage, frequency tables, mean and the like. In addition, ANOVA or analysis of variance will be used to compare

| |
|---|
| differences regarding socioeconomic impacts for the three indicator variables along with its items, such as demographic indicator, access to social infer structure and economic indicators due to water hyacinth infestation. A diagnostics test will be done before ANOVA to check whether or not the data would normally have distributed using Shapiro-Wilk Test. The values of Shapiro-Wilk Test for both variables will be found to be above 0.05. This indicates that data would not be deviated from a normal distribution. Homoscedasticity test will be done and the value of the test has to be above 0.05. This will improve the homogeny of the data variance. At all stages, a regression analysis will be employed to assess the strength of the relationship between variables so as to use for modelling the direct and indirect impacts. Also, the regression helps to apply a model for multiple independent variables, and to assess interaction terms to determine the effect of one independent variable which depends on the value of another variable. Since, a quantitative data is a numerical observation and as mentioned above the phenomenon will be analysed by descriptive statistics using SPSS 26.0 statistic software for Windows, 2013. Likewise, the qualitative data collected will be characterized by more of descriptive analysis aimed at creating a common understanding of the subject being studied. GIS and remote sensing will be used throughout the analysis for triangulation, which can be explained by using the tools of analysis often designed for GIS and remote sensing image and pictures. The audio-records that will be taken during FGDs and KIIs will be first transcribed into English and systematically read and lastly, transcripts will be analysed using thematic analysis. In the analyses the results can quantify current and potential infestations, and the findings will inform the better management strategy for the study area and beyond. |

| | |
|---|---|
| **5.2** | **Provide the contact details of the statistician or external coder that you will use** (if applicable) |
| NA | |

| | |
|---|---|
| **5.3** | **For a quantitative study or phase of your research, provide a brief description of the measures YOU WILL TAKE with regard to your study to ensure validity and reliability, taking into account:** |
| **(a) Internal and External validity of the research design**<br>This research design can be reproducible under similar concomitance when repeated and checking the consistency of results across different observers, across time, and across parts of the test itself is possible. For example, mapping the water hyacinth expansion or water sample laboratory test in triplicate and questionnaire prepared to be administered for respondents have filly filled the criteria of both Reliability and validity<br><br>**(b) Validity and Reliability of data gathering instrument**<br><br>As long as a valid measurement is reliable an item in this research can produces an accurate result which can be reproducible and, in this case, it is possible to prove the measurements consistency. | |

| | |
|---|---|
| **5.4** | **For a qualitative study or phase of your research, provide a brief description of the measures YOU WILL TAKE with regard to your study to ensure trustworthiness and/or authenticity, for instance taking into account:** |

a) **Credibility: all the secondary data will be collected from accredited governmental organizations and also for primary data collection a set of scientifically proven methodologies will be followed.**

b) **Dependability: Since the research is totally for academic purpose there will be no possible influence in the research process or in research findings. It is completely independent.**

c) **Conformability; it is obvious and real that in Ethiopia water hyacinth expansion is becoming an issue causing serious problem in the Environment and in the biodiversity.**

d) **Transferability; this research has a high degree to which the results can be transferred to other contexts or settings when ever it is important to transfer.**

e) **Authenticity: the finding of this research will have true meaning if properly the study will have been conducted area and its finding will add a real value to the locality in one or another way.**

| **5.5** | **List the references used in the application form** |
|---|---|
| References: 1. Munyaradzi Chitakira<br>2. Luxon Nhamo | |
| **5.6** | **Indicate the timeline** *(*Insert additional rows if necessary)* |
| **Planned research activities** **(i.e. ethics clearance, data collection, analysis, report writing, editing, printing, etc.)** | **Anticipated completion time** |
| Ethics clearance, | End of July 2021 |
| Data collection, | End of August up to End of October 2021 |
| Data analysis, | Start of November up to End of December 2021 |
| Report writing, | After December 2021 up to March 2022 |
| Editing, | March 2022 up to May 2022 |
| Printing, etc.) | Tentatively from End of May up to July 2022 |
| **5.7** | **Indicate the budget to** *justify the financial feasibility of the research (*Insert additional rows if necessary)* |
| **Planned research activities** | **Estimated cost in SA Rand** |
| Research activities | 100,000.00 |
| Additional Support (travelling, conference, and others, (no accommodation nor compensation & meals) | 20,000.00 |
| Total | **120,000.00** |
| **5.8** | **If you do run out of funding to cover your budget, do you have a contingency fund or additional** |

#### SECTION 6: ETHICAL CONSIDERATIONS

| **6.1 (a)** | **Describe the process of obtaining Informed Consent below** |
|---|---|

**PARTICIPANT INFORMATION SHEET**

Ethics clearance reference number: 2021/CAES_HREC/090
Research permission reference number: REC-170616-051

Date: 6/28/2021

Title: **Investigating current and potential expansion, socioeconomic impact and climate smart management approach of water hyacinth (Eichhornia crassipes) in the rift valley lakes of Ethiopia**.

**Dear Prospective Participant**

My name is Esayas Elias Churko and I am doing research with Dr L Nhamo, a senior lecturer in the Department of Environmental Science towards a PhD at the University of South Africa. We have funding no funding except students Bursary from UNIISA. We are inviting you to participate in a study entitled: Investigating current and potential expansion, socioeconomic impact and climate smart management approach of water hyacinth (Eichhornia crassipes) in the rift valley lakes of Ethiopia. I am conducting this research to find out the current and potential expansion, socioeconomic impact and climate smart management approach of water hyacinth (Eichhornia crassipes) in the rift valley lakes of Ethiopia. This study is expected to collect important information that could describe current and potential expansion, its socioeconomic impact and climate smart management approach of water hyacinth (Eichhornia crassipes) to the community and beyond. You have chosen from this locality because you are one the residents live at nearby distance to the lakes, and the information gathered here will remain confidential and legal as to *the Protection of Personal Information Act, nr 4 of 2013. Just to let you know in this research*, *including you,* approximately *400 respondents would participate.* Your actual role in ihe study involves responding *questionnaires* in this document in total which takes 25 minutes. Your participation is voluntary, has no payment nor that there is no penalty or loss of benefit for non-participation. You are free to withdraw at any time and without giving a reason and it has no negative consequence in your life. *A report of the study may be submitted for publication, but individual participants will not be identifiable in such a report*. Hard copies of your answers will be stored by the researcher for a period of five years in a locked cupboard/filing cabinet *[where? Indicate the location]* for future research or academic purposes; electronic information will be stored on a password protected computer. This study has received written approval from the Health Research Ethics Committee of the College of Agriculture and Environmental Sciences, Unisa. Should you have concerns about the way in which the research has been conducted, you may contact:

Supervisors contact details hereby below:

Dr L Nhamo

; 073-166-5859
Prof M Chitakira
; 011-471-3220

The research ethics chairperson of the CAES Health Research Ethics Committee,

Prof MA Antwi

on 011-670-9391 (if you have any ethical concerns)

Thank you for taking time to read this information sheet and for participating in this study.

Thank you.

X

Esayas Elias Churko
PhD Researcher

Esayas Elias Churko

**(b)**

Insert the information sheet and informed consent document(s) here. Note: *The participant information sheet ought to explain all criteria stipulated below in 6.2, with the exception of those marked with **

#### CONSENT TO PARTICIPATE IN THIS STUDY

I, ____________________________ (participant name), confirm that the person asking my consent to take part in this research has told me about the nature, procedure, potential benefits and anticipated inconvenience of participation.

I have read (or had explained to me) and understood the study as explained in the information sheet.

I have had sufficient opportunity to ask questions and am prepared to participate in the study.

I understand that my participation is voluntary and that I am free to withdraw at any time without penalty (if applicable).

I am aware that the findings of this study will be processed into a research report, journal publications and/or conference proceedings, but that my participation will be kept confidential unless otherwise specified.

I agree to the recording of the <insert specific data collection method>.

I have received a signed copy of the informed consent agreement.

Participant Name & Surname………………………………………

Participant Signature………………………………………Date…………………

Researcher's Name & Surname………**Esayas Elias Churko….**

Researcher's signature…………………………………...Date…………………

| 6.2 | **Checklist to ensure that the <u>participant information sheet</u> and <u>consent form</u> meet ethical requirements**<br>***Standard participant information sheets and consent forms are available on the research website or can be requested from*** | **YES** | **NO** | **N/A** |
|---|---|---|---|---|
| | *Place an 'x' in the box provided* | | | |
| a) The identity and position of the researcher(s) and the organisation collecting the information? | | **'x'** | | |
| b) The purposes for which the information is being collected? | | **'x'** | | |
| c) Reason why the participant has been selected and procedures for selecting participants? | | **'x'** | | |
| d) Participant's actual role in the study? | | **'x'** | | |
| e) Expected duration of participation? | | | | **'x'** |

| | | | |
|---|---|---|---|
| f) Statement that participation is voluntary and that there is no penalty or loss of benefit for non-participation? | | | **‘x’** |
| g) Benefits to the participant and others? | | **‘x’** | |
| h) Potential risks <u>as well as</u> measures that will be taken if injury or harm attributable to the study occurs? | **‘x’** | | |
| i) Statement that participant can withdraw at any time without obligation to explain or any adverse effects? | **‘x’** | | |
| j) Compensation/gifts/services for participants? | **‘x’** | | |
| k) Reimbursement and any costs incurred by participants? | | | **‘x’** |
| l) Indemnity if applicable? * | | | **‘x’** |
| m) The period for which the records relating to the participant will be kept? | **‘x’** | | |
| n) The steps taken to ensure confidentiality and secure storage of data? | **‘x’** | | |
| o) The types of individual or organisation to which your organisation usually discloses information of this kind? | **‘x’** | | |
| p) How privacy will be protected in any publication of the information? | **‘x’** | | |
| q) How feedback will be provided? | **‘x’** | | |
| r) Any exclusion to confidentiality? (e.g. when focus groups are used) * | | | **‘x’** |
| s) Consent for third party sharing of data e.g. statistician, coders if applicable* | | | **‘x’** |
| t) Consent for cross border data transfer if applicable* | | | **‘x’** |
| u) Consent for data sharing in a credible public repository if applicable* | | | **‘x’** |
| v) The steps taken to ensure that cultural protocol has been observed* | | | **‘x’** |

| **6.3** | **Checklist to ensure that the process of <u>obtaining assent</u> meets ethical requirements (IF APPLICABLE).** | **YES** | **NO** |
|---|---|---|---|
| *Not applicable* [ ] <br> *Place an ‘x’ in the box provided* | | | |
| a) A statement of the purpose of the research or study? | | X | |
| b) A description of the procedure to be applied in dealings with the minor? | | X | |
| c) A statement that the minor’s identity will not be revealed? | | X | |
| d) A description of the potential risks or discomfort associated with the research? | | | |
| e) A description of any direct benefits to the minor? | | | X |
| f) A statement that the minor is not compelled to participate? | | | X |
| g) A statement that the minor is free to withdraw at any time? | | X | |
| h) A statement that the minor should discuss participation with the parents prior to signing the form? | | X | |
| i) A statement that the parent(s)/guardian(s) of the minor will be asked for permission on behalf of the minor? | | X | |
| j) A statement that the parent(s)/guardian(s) of the minor will receive a copy of the signed form? | | X | |
| k) Invitation to ask questions? | | X | |
| l) Contact details of researcher? | | X | |
| Note that only the minor and the researcher obtaining assent should sign the child assent form. A copy of the child assent form should be given to the parent or legal guardian. | | | |

*[*If other, provide details in the space allowed for comments]*

| **6.4** | **Measures taken to protect confidentiality:** <br> *(adapted from the University of Cape Town, research ethics application form)* |
|---|---|

| | | |
|---|---|---|
| 6.4.1 | Paper-based records must be kept in a secure location and should only be accessible to personnel involved in the study.<br>**Please indicate who will have access to the data and where will it be retained.** | |
| The PhD researcher or doctoral student who will have access to the data and be retained. | | |
| 6.4.2 | Computer-based records must only be available to personnel involved in the study through the use of access privileges and passwords.<br>**Please indicate who will have access to the computer-based records.** | |
| The PhD researcher or doctoral student who will have access to the data. | | |
| 6.4.3 | Personnel will be required to sign statements agreeing to protect the security and confidentiality of identifiable information.<br>**Please indicate who will be required to sign confidentiality agreements and append the agreements to the application.** | |
| The PhD researcher or doctoral student who will have access to the data and where it will be retained. | | |
| 6.4.4 | *Place an 'x' in the box provided* | |
| i | Personal identifiers will be removed from research-related information | X |
| ii | Encryption | X |
| iii | Use of pseudonyms | X |
| iv | Participants in focus groups will be advised that confidentiality cannot be assured | X |
| Comments: the points of for discussion with the team of FGD does not require high level of confidentiality even if the group does not assured confidentiality. | | |

| | |
|---|---|
| **6.5** | **Data sharing (if applicable)**<br>*In the South African context, agencies including the National Research Foundation (NRF) require that data supporting publications be deposited in an accredited open access (OA) Data repository from March 2015 onwards with a registered Digital Object Identifier (DOI) for citation and referencing.* |
| 6.5.1 | How will you obtain consent from the research participants and/or other stakeholders i.e. funders, participating institutions, etc. for further use of research information? Note that personal identifiable data cannot be shared without explicit prior consent. |
| The Request letter will be offered officially forwarded from the UNISA centre Ethiopia, for all the respective bodies in their physical address or by email. | |
| 6.5.2 | Where, with whom and how will you share the data?<br>With no one I do share the data, except assigned academic advisors at UNISA |
| 6.5.3 | How will you de-identify data to protect participants' privacy during further processing? |
| Removing names and ID number or telephone from the consent paper or page. | |

| | |
|---|---|
| **6.6** | **Data Storage and Procedures for Disposal of the data** |
| 6.6.1 | For what period of time will the data be retained? The Unisa Policy on Research Ethics stipulate that data should be retained for a minimum period of 5 years. *Please note that this time period presents a minimum standard.* |
| Yes, it presents the standard. | |
| 6.6.2 | What reasonable steps will be taken to dispose of or permanently de-identify personal information if it is no longer needed for the purpose of research? |
| Hiding names and any other personal details is possible in this research. | |

| | |
|---|---|
| **6.7** | **How will participants be informed of the findings or results and consulted on potential or actual benefits of such findings or results to them or others?** (Copy of journal article, book, chapter, summary report to organisation, on-line web based, oral presentation, other) |
| At the end of publication, it is possible to den copy of journal article, book, chapter, summary report to organisation, on-line web based, oral presentation, other to the offices. | |
| **6.8** | **In the case of the use of indigenous knowledge, how will you ensure that participants are not exploited/harmed for instance by protecting their Intellectual Property rights?** (if applicable) |
| NA | |

| | |
|---|---|
| **6.9** | **How will participants benefit, gain access or share in products developed from the study in case of collaborative projects or the use of indigenous knowledge?** (if applicable) |
| NA | |

| | |
|---|---|
| **6.10** | **Indicate how you envisage publishing this research.** (thesis, journal article, book, chapter, on-line web based, oral presentation, other) |
| For publishing this research, it will be handy to classify each article based on specific objective. | |

| | |
|---|---|
| **6.11** | **Describe the nature and amount of compensation including reimbursements, gifts, services or incentives to be provided to each group of participants.** (if applicable) |
| In total amount of compensation including reimbursements, gifts, services or incentives to be provided to each group of participants will be per dime and it will be about 400 SA Rand/ day. | |
| **6.12** | **Describe any financial costs that might be incurred by participants. If participants incur costs, how will you ensure that it is fair?** (if applicable) |
| NA | |

**PLEASE REMEMBER TO COMPLETE AND APPEND THE CHECKLIST TO YOUR APPLICATION – APPENDIX 1**
